# Neuronopathic GBA1 L444P mutation accelerates Glucosylsphingosine levels and formation of hippocampal alpha-synuclein inclusions

**DOI:** 10.1101/2022.04.07.487391

**Authors:** Casey L. Mahoney-Crane, Megha Viswanathan, Dreson Russell, Rachel A.C. Curtiss, Jennifer Freire, Sai Sumedha Bobba, Sean D. Coyle, Monika Kandebo, Lihang Yao, Bang-Lin Wan, Nathan G. Hatcher, Sean M. Smith, Jacob N. Marcus, Laura A. Volpicelli-Daley

## Abstract

The most common genetic risk factor for Parkinson’s disease (PD) is heterozygous mutations in the GBA1 gene which encodes for the lysosomal enzyme, glucocerebrosidase (GCase). GCase impairments are associated with an accumulation of abnormal α-synuclein (α-syn) called Lewy pathology, which characterizes PD. PD patients heterozygous for the GBA1 L444P mutation (GBA1^+/L444P^) have a 5.6-fold increased risk of cognitive impairments. In this study, we used GBA1^+/L444P^ mice to determine the effects of this severe GBA1 mutation on lipid metabolism, expression of synaptic proteins, behavior, and α-syn inclusion formation. GBA1^+/L444P^ mice showed reduced GCase activity in limbic brain regions and expressed lower levels of hippocampal vGLUT1 compared to wildtype (GBA1^+/+^) mice. GBA^+/L444P^ mice also demonstrated impaired fear conditioning, but no motor deficits. We show, using mass spectrometry, that mutant GCase and age increased levels of glucosylsphingosine (GlcSph), but not glucosylceramide (GlcCer), in the brains and serum of GBA1^+/L444P^ mice. Aged GBA1^+/+^ mice also showed increased levels of GlcSph, and decreased GlcCer. To model disease pathology, templated α-syn pathology was used. α-Syn inclusions were increased in the hippocampus of GBA1^+/L444P^ mice compared to GBA1^+/+^ mice, but not in the cortex, or substantia nigra pars compacta (SNc). Pathologic α-syn did not cause a loss of dopamine neurons in the SNc. Treatment with a GlcCer synthase inhibitor prevented loss of cortical α-syn inclusions, but not loss of dopamine neurons. Overall, these data suggest the critical importance to evaluate the contribution of hippocampal pathologic α-syn and brain and serum glucosylsphingosine in synucleinopathies.

**SIGNIFICANCE STATEMENT:** Synucleinopathies, such as Parkinson’s disease (PD) and Dementia with Lewy bodies (DLB), are both pathologically characterized by abnormal α-synuclein (α-syn). Mutant GBA1 is a risk factor for both PD and DLB where a reduction of glucocerebrosidase (GCase) activity is seen. Collectively, this indicates the significance of evaluating mutant GCase in synucleinopathies. Our data suggest the critical importance to evaluate the contribution of hippocampal pathologic α-syn and brain and serum glucosylsphingosine (GlcSph) accumulation in synucleinopathies. Moreover, these pathologic outcomes may contribute to the nonmotor symptoms clinically observed in PD and DLB. Our findings highlight the importance of GlcSph as a relevant biomarker for future therapeutics.

## INTRODUCTION

Synucleinopathies, such as Parkinson’s disease (PD) and Lewy body dementias, are characterized by the neuronal accumulation of α-synuclein (α-syn) aggregates which are phosphorylated (p-α-syn), called Lewy pathology. Historically, most research has focused on the role of Lewy pathology in the death of dopamine neurons because of the contribution to motor symptoms of PD. However, Lewy pathology is significantly correlated with increased Parkinson’s disease dementia (PDD) (Irwin et al., 2012). PD patients with the severe L444P mutation in the GBA1 gene show up to a 5.6-fold increase in the risk for dementia (Cilia et al., 2016). The severity of the GBA1 L444P mutation can be classified based on the location of the mutation contributing to unstable glucocerebrosidase (GCase) due to a conformational change (Dvir et al., 2003; Migdalska-Richards and Schapira, 2016). In addition, the GBA1 L444P heterozygous mutation is one of the most prevalent mutations in PD. Nearly 8-12% of PD cases occur due to a mutation in GBA1 (GBA1-PD), which is the most common risk factor for PD, after aging (Stoker and Greenland, 2018). The severe GBA1^+/L444P^ mutation represents approximately ∼35% of all GBA1 mutations linked to PD (Sidransky et al., 2009). Thus, the interaction between mutant GBA1 and the formation of Lewy pathology may contribute to cognitive decline in PD.

*GBA1* encodes for the lysosomal enzyme, GCase, which metabolizes the sphingolipids glucosylceramide (GlcCer) and glucosylsphingosine (GlcSph) to glucose and ceramide or sphingosine (Do et al., 2019; Boer et al., 2020). The GBA1 L444P mutation impairs proper folding of the enzyme, preventing its transport from the endoplasmic reticulum to the Golgi apparatus and lysosomes. Previous studies evaluating the heterozygosity of GBA1 L444P revealed a reduction of GCase activity by approximately 40% (Fishbein et al., 2014). Models of α-synucleinopathies show that mice heterozygous for GBA1 L444P have increased pathologic α-syn aggregate formation and motor defects (Taguchi et al., 2017; Migdalska-richards et al., 2020), suggesting reduced GCase activity promotes accumulation of Lewy pathology.

Here, we show reduced GCase activity in the cortex, striatum, hippocampus, and midbrain of GBA1^+/L444P^ knock-in mice relative to wildtype (GBA1^+/+^) mice. GBA1^+/L444P^ mice and aged GBA1^+/+^ mice show increased brain levels of glucosylsphingosine, but not glucosylceramide which is consistent with previous findings (Taguchi et al., 2016; Polinski et al., 2021). GBA1^+/L444P^ mice also show reduced levels of the presynaptic protein, vGLUT1, and impairments in fear conditioning, but not motor behavior, suggesting that mutant GBA1 alone causes neuronal defects in limbic brain regions. To determine the impact of GBA1^+/L444P^ on the formation of α-syn aggregates and neuronal toxicity, mice were analyzed after injections of α-syn pre-formed fibrils to induce corruption of endogenously expressed α-syn. There was increased α-syn pathology in the hippocampus of GBA1^+/L444P^ knock-in mice compared to GBA1^+/+^ mice, but this did not occur in the substantia nigra pars compacta (SNc) or cortex. Expression of GBA1^+/L444P^ did not enhance α-syn inclusion-induced loss of dopamine neurons in the SNc compared to GBA1^+/+^ mice. Treatments with a glucosylceramide synthase inhibitor did not prevent α-syn inclusion formation in primary cultured neurons or in fibril-injected mice. Overall, these data suggest that expression of GBA1 L444P 1) increases the susceptibility of hippocampal neurons to dysfunction, 2) increases the formation of α-syn inclusions in the hippocampus, and 3) supports findings that GlcSph is a highly salient biomarker for the effects of GBA1 PD therapeutics, more so than GlcCer.

## MATERIALS AND METHODS

### Animal care

All animal protocols were approved by the University of Alabama at Birmingham’s Institutional Animal Care and Use Committee. Both male and female GBA1 mice heterozygous for the L444P mutation (GBA1^+/L444P^), obtained from the Mutant Mouse Resource and Research Center (MMRRC) at the University of North Carolina at Chapel Hill, were used for these studies and generated as previously described (Liu et al., 1998). Mice were on a 12-hour light/dark cycle and had *ad libitum* access to food and water.

### Fibril preparation

Fibrils were prepared as previously described (Stoyka et al., 2020, 2021). Mouse monomeric α-synuclein (α-syn) was purified from *E. coli*, and endotoxin was removed using the Pierce high-capacity endotoxin spin columns and quantified using the Pierce endotoxin chromogenic quantification kit (Volpicelli-Daley et al., 2014; Stoyka et al., 2021). Monomeric α-syn was stored at concentrations >10 mg/mL at −80°C. To generate fibrils, monomer was rapidly thawed and centrifuged at 20,000 *g*. The concentration of the supernatant was determined using A280 nm with an extinction coefficient of 7450 M^-1^ cm^-1^ using a NanoDrop 2000/2000c Spectrophotometer (ThermoFisher Cat no. ND-2000) and diluted to 5 mg/mL in 150 mM KCl and 50 mM Tris-HCl. A volume of 500 μL was shaken at 37 °C at 700 rpm for 7 days. The fibrils were spun for 10 min at 16,363 *g* at room temperature and resuspended in 250 μL of the above buffer. To determine the concentration of preformed fibrils (PFFs), 5μL was incubated with 5 μL of 8 M guanidinium chloride to dissociate the fibrils, and the concentration was measured using A280 nm. Buffer was added to the fibrils to obtain a final concentration of 5 mg/mL, aliquoted to 25 μL, and stored at −80 °C until use. On the day of injection, fibrils were sonicated using a QSonica700 cup horn sonicator with a 16°C water bath for 15 minutes at 30% amplitude for 3-seconds on and 2-seconds off pulse at 85,824 joules. Fragmentation of fibrils was confirmed using dynamic light scattering.

### Bilateral Striatal Injection of Fibrils

Three-month-old mice were anesthetized with vaporized isoflurane (1%) on a stereotactic set-up (KOPF, Model 940, small animal stereotaxic instrument, standard accessories). The scalp was shaved and wiped with chlorhexidine, and a vertical incision was made. Bilateral coordinates relative to bregma were +1 mm anterior, ± 2.0 mm lateral, and −3.2 mm ventral relative to the skull. Two μL of sonicated fibrils (5 µg/µL) or monomeric α-syn (5 µg/µL) was injected at a constant rate of 0.5 μL/min. The needle was left in place for four minutes to allow diffusion, then retracted 1 mm, then left in place for another one minute before fully retracting the needle. Scalp incisions were closed with EZ-Clips. Mice were administered 10 units of 1.5mg/mL carprofen subcutaneously the next day.

Starting one-week post-injections, mice were fed chow containing venglustat or control chow without venglustat (Research Diets) provided by Merck & Co., Inc., Kenilworth, NJ, USA. Male and female mice were fed separate chow formulations with the concentration of venglustat targeting a dose of 60 mg venglustat/kg body weight/day based on the mean expected food consumption and the mean body weight of male and female mice. Cages were allocated 5g chow per mouse, and food was replenished every week, or as needed. Mice were kept on the formulated chow until the day of sacrifice.

### Behavior testing

Behavioral testing was performed as previously described (Stoyka et al., 2020, 2021). Each test was run with operator silence and, when available, the assistance of a white noise machine to minimize potentially intrusive noises. The operator was blinded to treatment and genotype.

### Open Field

Open Field test was run on each mouse using ActivityMDB program to track motion in a 1×1-foot enclosed field with two zones of detection: zone 0 and zone 1. Zone 1 was defined as the center of the box, whereas zone 0 was the perimeter of the box. This tracking monitored the velocity of movement, and time spent in each zone over the course of 10 minutes. The percent time in the center was calculated as (the time spent in zone 1 divided by the total time spent in the box) *100.

### Pole Test

The pole test was conducted using a 12-inch wooden dowel rod with a diameter of 1.27 cm, anchored with a 4×6×1-inch block of wood and ridged at 1-inch intervals with rolled rubber bands. The dowel rod was placed vertically in the center of a 1×1 foot enclosure. The mice were trained once by being placed face-down at the base of the pole and allowed to climb off onto the floor of the enclosure before being returned to their cage. Following this, each mouse was placed face-up at the top of the pole and allowed no more than 180 seconds to turn around and climb down to the bottom; this was repeated five times. Mice were given a minimum of 30 seconds to rest between each trial. Three time point measurements were recorded. The first time point started from the time the mouse was placed on the pole to the time taken for the mouse to reach the bottom (total descent time). Measurements were recorded for the time the mouse took to turn around on the pole to face the bottom (turnaround time) and the total time to descend (time descending) in seconds. The average time for each time point was calculated from all five trials for each mouse.

### Fear conditioning

Fear Conditioning was performed using an apparatus designed with a grid of metal bars on the floor and connected to an electrical output box set to 0.6 mAmp controlled by the VideoFreeze software. On day 1, mice were individually placed in the apparatus and, after two minutes, a 30-second tone was presented with a two-second mild-foot shock at the last two seconds of the tone. The mouse was allowed to rest for 60 seconds, followed by a second delivery of tone and shock. The mouse was allowed to rest for 60 seconds after the second mild foot shock. This was completed once per mouse. Freezing was measured manually by a researcher blinded to treatment and genotype. Percent time freezing was calculated as the time (in seconds) the mouse did not move divided by the total time of each time point (e.g., 0-120s or 120-150s). The conditioned stimulus (CS) was defined as the tone played, and the unconditioned stimulus (US) is the mild foot shock.

The following day, the mice performed the “Contextual” test. The mice were placed in the fear conditioning apparatus with an identical environment as training. Freeze times of each mouse over the course of 5 minutes (300 seconds) were recorded. Percent time freezing was calculated as the time (in seconds) the mouse did not move divided by the total time. The first minute (60s) was graphed.

On day three, “Cued” fear conditioning was performed in the same apparatus by replacing the metal bars on the bottom with solid white tiles. Solid white tiles were also placed inside the monitored area, changing the environment from a square cage with metal bars on the floor to a solid white triangular container. The cage was cleaned using 70% Ethanol and treated with a small drop of concentrated peppermint oil to conceal any familiar scents. The area surrounding the cage was changed by placing a purple and white striped pattern along the wall. The same CS used for training was presented for 30 sec, without the US, followed by a 60-sec rest and repeated twice for a total of 300 sec. Freeze times of each mouse were measured over the course of 5 minutes (300 Seconds). The average percent time the mouse spent freezing during the two tones was graphed.

### Immunohistochemistry (IHC) and Immunofluorescence (IF)

Ten months post-injection, mice were anesthetized with isoflurane and transcardially perfused with saline solution (0.9% NaCl, 0.005% sodium nitroprusside, and 10 units/ml heparin sodium) followed by cold 4% paraformaldehyde (PFA) in phosphate-buffered saline (PBS) using gravity perfusion. Brains were removed, and postfixed for 24 hours in 4% PFA followed by 30% sucrose PBS solution for 48 hours. Brains were then frozen in methylbutane solution (−50 to −60 °C) and sectioned into 40 μm serial sections with a freezing microtome (Leica SM 2010R). Serial sections spaced 240 μm apart representing the entire forebrain were collected in a 6-well tray using a freezing microtome (Stoyka et al., 2020, 2021). Sections were stored at −20 °C in a solution of 50% glycerol, and 0.01% sodium azide in Tris-buffered saline (TBS).

All incubations and rinses were performed using an orbital shaker. For IHC, sections were rinsed three times in TBS and quenched in 3% H_2_O_2_ in TBS for 10 min. Following three rinses in TBS, sections were incubated in antigen retrieval solution (10 mM sodium citrate, 0.05% Tween 20, pH 6) at 37°C for 1 h, rinsed three times in TBS, and blocked using 5% normal donkey serum (Equitech-Bio Inc) with 0.1% Triton X-100 in TBS for 1 h at 4°C. After blocking, sections were incubated in primary antibody solution of rabbit polyclonal antibody to tyrosine hydroxylase (TH; Sigma AB152, RRID: AB_390204) in 5% normal donkey serum in TBS at 4°C overnight. After three rinses, sections were then incubated in Biotin-SP AffiniPure Donkey Anti-Rabbit IgG H&L (Jackson ImmunoResearch 711-065-152) in 5% normal donkey serum in TBS for 2 h, followed by incubation with Avidin-Biotin Complex Peroxidase Standard Staining kit reagent (ABC; Thermo Scientific) for 1 h at room temperature. Sections were developed using ImmPACT-3 3’-diaminobenzidine (DAB; Vector Labs), sequentially dehydrated as previously described (Stoyka et al., 2020, 2021), and mounted on charged slides using Permount.

For IF, sections were rinsed three times in TBS and incubated in an antigen retrieval solution (10 mM sodium citrate, 0.05% Tween-20, pH 6.0) for 1 hour at 37 °C. Sections were rinsed three times in TBS and blocked and permeabilized in 5% normal goat serum, 0.1% TritonX-100 in TBS for 1 hour at 4 °C. Sections were rinsed three times in TBS and then incubated in a primary antibody solution with 5% normal goat serum in TBS overnight at 4 °C. Primary antibodies used included phosphorylated α-syn (Abcam, Cat#ab51253, RRID:AB_869973), TH (Abcam, Cat#ab76442, RRID:AB_1524535), and total synuclein (BDTrans, Cat#610786, RRID:AB_398107). Sections were rinsed three times in TBS and incubated with secondary antibodies (1:500) in 5% goat serum in TBS overnight at 4 °C. Secondary antibodies were AlexaFluor goat anti-rabbit 555 IgG (ThermoFisher, Cat#A-21429, RRID:AB_2535850), AlexaFluor goat anti-mouse 647 IgG_1_ (ThermoFisher, Cat#A-21240, RRID:AB_2535809), AlexaFluor goat anti-chicken 488 IgY (ThermoFisher, Cat#A-11039, RRID:AB_2534096), and Hoechst 33342 (1:1000). Sections were rinsed three times in TBS and mounted on charged glass slides with Prolong Gold (Life Technologies).

### Quantification of p-α-syn Inclusions in Brain Sections

Images and quantitation were performed by a researcher blinded to experimental conditions. Sections were imaged on Nikon C2 confocal microscope at 10X magnification and analyzed using ImageJ software. Images of the basolateral amygdala, hippocampus, SNc, and dorsomedial prefrontal cortex (dmPFC) were captured from the fibrils-injected brains. Representative images were matched across samples to ensure consistency in the relative positions (according to Bregma coordinates) of the regions imaged. One image from one of the hemispheres was captured of the dmPFC and one image each from both hemispheres was captured for the other regions of interest. Several images were tiled together to create a single, large-field image of the larger regions, the dmPFC, and hippocampus, using the Nikon Elements imaging software.

p-α-Syn inclusions were quantified using the CellCounter program on ImageJ. Regions of interest were drawn manually on ImageJ. For the hippocampal images, separate regions of interest (dentate gyrus, CA1, CA3) were drawn demarcating the layers. Images were thresholded using the “maximum threshold” option on FIJI, by setting the same maximum brightness for all images for each region. The threshold was set to 180 for hippocampal and SNc images, 810 for all amygdala images, and 315 for all dmPFC images. For entorhinal cortical images, images were not thresholded. This ensured that all inclusions were clearly visible. Neuritic and somal aggregates were counted manually using the CellCounter program, and each hemisphere was evaluated separately. The average of both hemispheres for each mouse was calculated and graphed.

### Unbiased Stereology

Unbiased stereology was conducted using the Olympus BX51 bright field microscope and the optical fractionator probe (Stereo Investigator software, Stereology Resource Center) to analyze TH neurons in the SNc. On average, five to six sections of the SNc from rostral to caudal (spaced 160 µm apart) were quantified. Quantification was conducted with a counting frame of 50um by 50um, optical dissector height of 22 um, and section thickness of 33um. The measured counting variability for Schmitz-Hof coefficient of error (CE) was below 0.135 for all mice.

### Primary hippocampal neurons

Embryonic day 16-18 embryos from a GBA1^+/L444P^ cross were removed and hippocampal tissue from each embryo was dissected and placed in individual falcon tubes containing 10mL Hibernate E solution. Neurons were cultured as previously described (Volpicelli-Daley et al., 2014) and neurons from each embryo were plated separately in 24-well plates with coverslips. Tails from each embryo were used to determine genotype.

### Treating primary hippocampal neurons

On day in vitro (DIV) 7, fibrils were diluted to 100 μg/mL in PBS and sonicated using a probe-tip sonicator (Fisher Scientific, Model CL-18) for 30 seconds with a 1-second pulse at 30% amplitude. Sonicated fibrils were diluted to 0.1 μg/mL in neuronal media. Neurons were treated with fibrils (0.1 μg/mL) or monomers (0.1 μg/mL) and 100 nM Eliglustat (Sigma) or equal volume DMSO.

### Immunofluorescence of Cultured Hippocampal Neurons

Neurons were fixed in 4% paraformaldehyde/4%sucrose solution in PBS at room temperature for 30 minutes. After rinsing four times in PBS, neurons were blocked and permeabilized with 3%BSA and 0.05% saponin in PBS for 30 minutes at room temperature. Neurons were then incubated in primary antibody in 3% BSA and 0.05% saponin overnight at 4°C. Primary antibodies included were Neurofilament heavy polypeptide (Abcam, Cat#ab722996, RRID: AB_2119618), phosphorylated α-syn (Abcam, Cat#ab51253, RRID: AB_869973), and Total α/β-synuclein 202 (BioLegend, Cat#836602, RRID: AB_2734606). Neurons were rinsed three times in PBS with 0.05 % saponin and incubated in secondary antibody in 3%BSA and 0.05% saponin for 1 hour at room temperature. Secondary antibodies included were AlexaFluor goat anti-rabbit 488 IgG (ThermoFisher, Cat#A-11034, RRID: AB_2576217), AlexaFluor goat anti-mouse 555 IgG_2a_ (ThermoFisher, Cat#A-21137, RRID: AB_2535776), AlexaFluor goat anti-chicken 647 IgY (ThermoFisher, Cat#A-21449, RRID: AB_2535866), and Hoechst 33342 (1:1000). Neurons were then rinsed twice in PBS with 0.05 % saponin and three times with PBS and mounted with Prolong Gold (Invitrogen). Five representative fields per coverslip were imaged using Nikon A1R confocal microscope with a 40X oil-immersion objective. Images were analyzed using ImageJ software by selecting the “IsoData” auto-threshold option. Percent area occupied by fluorescent signal was calculated using “particle analysis” in ImageJ. Percent area of p-α-syn was normalized by dividing by the percent area occupied by neurofilament-H immunofluorescence.

### DQ-BSA Red in Cultured Hippocampal Neurons

DQ-BSA Red (Invitrogen) was solubilized in 1 mL PBS to a stock concentration of 1mg/mL and sonicated with a probe tip sonicator for 5 seconds with 1-second pulse at 20% amplitude and then passed through a 1 mL syringe with a 0.22 μm filter (Millipore Sigma). Neurons were incubated with 25 μM of DQ-BSA Red solution diluted into neuronal media for 2 hours at 37°C. After DQ-BSA Red treatment, neurons were fixed in 4% paraformaldehyde/4%sucrose solution in PBS at room temperature for 30 minutes. Neurons were then rinsed three times with PBS and incubated in Hoechst 33342 for 30 minutes at room temperature. Neurons were rinsed twice in PBS and mounted with Prolong Gold (Invitrogen). Coverslips were imaged on Zeiss Axiovert.Z1 widefield microscope at 63X oil objective.

### DQ-BSA Red Analysis

In ImageJ, Regions of Interest (ROIs) were drawn around the cell bodies. Hoechst 33342 staining was used as a guide to eliminate unhealthy or dead cells. Five fields were imaged per coverslip. Within each field, the Corrected Total Cell Fluorescence (CTCF) of an average of 5-15 cells was quantified per field as described in Marwaha and Sharma, 2017 (Marwaha and Sharma, 2017). The mean CTCF of all replicates for each sample was graphed.

### Sphingolipid quantification

#### Lipid extraction

Extraction of sphingolipids in cells was performed as described previously (Cosden et al., 2021). Briefly, primary cortical neurons were provided as frozen monolayers in 96 well plates. Extraction was performed within sample wells by addition of 150 μL MeOH spiked with a mix of stable isotopically labeled synthetic standards (d3 glucosylceramide d18:1/16:0, ^13^C_6_ glucosylsphingosine (1-(beta-D-Glucosyl-1,2,3,4,5,6-13C6)-sphingosine; (Matreya LLC. State College PA), d7 ceramide d18:1/16:0, d7 sphingosine, d7 sphingosine-1-phosphate and d9 sphingomyelin d18:/18:0 (Avanti Polar Lipids, Inc, Alabaster) at a final concentration of 10 ng/mL. Plates were capped, shaken for ten minutes at room temperature, and centrifuged at 1500*g* for five minutes before the transfer of supernatants to new 96 well plates for mass spectrometry analyses.

Plasma sphingolipid extraction was performed using CHCl3:CH3OH (1:2) buffer spiked with 0.5ng/ml of the SIL lipid standard mix from above. 0.4 mL of extraction buffer was mixed with 20 μL of plasma, shaken for ten minutes at room temperature, and centrifuged at 3000g for ten minutes. 300 μL of supernatant was removed and dried down using N2 evaporation. Dried samples were reconstituted in 50 μL methanol immediately prior to analyses by mass spectrometry. Extraction of glycosphingolipids from brain homogenates was adapted from the method established previously (Hamler et al., 2017). Brain homogenates were prepared from pre-weighed frozen tissues sections by addition of MeoH:H2O (1:1) to generate an equivalent wet weight to volume ratio.

Homogenization was performed at 200 oscillations/min for two minutes using a QIAGEN TissueLyser II bead mill (QIAGEN Inc.-USA, Germantown, MD). ^13^C_6_ glucosylsphingosine and d5 glucosylceramide d18:1/18:0 (Avanti) internal standards were prepared as a DMSO stock solution at 2 μg/mL and 0.02 μg/mL, respectively. 25 μL of the internal standard stock solution was added to 100 μL of homogenate before the addition of 400 μL of acetone/MeOH buffer (1:3). Extracts were shaken for three minutes, resuspended with 100 μL of w,ater and centrifuged at 4000g for ten minutes. 500 µL of supernatant was removed, combined with 200 μL MeOH:H2O (1:1) and subjected to prefractionation via C18 solid phase extraction (SPE isolute C18, Biotage AB, Uppsala, Sweden). SPE eluates were evaporated to dryness under N2 gas and reconstituted in 50 μL DMSO and 200 μL mobile phase B liquid chromatography buffer (see below).

#### Mass spectrometry

Sphingolipids were quantified by liquid chromatography mass spectrometry using multiple reaction monitoring (MRM) scanning acquisition as described previously (Rocha et al. 2020). Separation was performed with a HALO HILIC 2.7 mm column (Advanced Materials Technology, Inc., Wilmington, DE) using a Waters Acquity UPLC (Waters Corp., Inc.). Mobile phase A and B consisted of 0.1% formic acid in H2O and 95% acetonitrile, 2.5% MeOH, 2.0% H2O, 0.5% formic acid and 5 mM ammonium formate, respectively. A ten-minute gradient was sufficient to achieve baseline separation of endogenous isomeric lipid species such as glucosylsphingosine and galactosylsphingosine. MRM acquisition was performed with a SciEx 5500 QTRAP mass spectrometer (SciEx LLC, Framingham, MA) operating in positive ion mode. Lipid quantification from primary cortical neurons was reported as ratios to internal standard and was used to compare relative changes in abundances across conditions. Absolute quantification was performed in plasma and brain tissue specimens using standard curves for each targeted species. For plasma lipid quantification, serial dilutions of a stock solution of 5 μg/mL of d5 GlcCer d18:1/18:0, ceramide d18:1/16:0, sphingosine d18:1, sphingosine-1-phosphate d18:1, sphingomyelin (porcine brain std; Avanti) and 50 ng/mL of glucosylsphingosine d18:1 (Avanti) were processed as plasma above, and standard curves were generated within each study plate. Absolute quantification of brain glucosyceramides was performed in tissue homogenates by spiking in serial dilutions down to 1/1000 of the d5 GlcCer d18:1/18:0 (1 mg/mL) stock solution for analysis as above. Glucosyslphingosine standard curves were generated using serial dilutions glucosylsphingosine d18:1 standard (Avanti) from DMSO stocks processed as above for LC-MS/MS analyses. Linear calibration curves with 1/x^2^ weighting factor were generated by plotting the ratio of the peak area of standards to internals standards. Total glucosylceramide was represented as the sum concentrations of C16:0 through C24:1 chain length variants.

### GCase activity assay

GBA1^+/+^ and GBA1^+/L444P^ mice were perfused with 30-40 mL of 0.9% NaCl saline (with 0.5% w/v sodium nitroprusside and 10 units/mL heparin) at 2 months of age. The brain was removed and dissected to isolate the midbrain, hippocampus, striatum, and cortex. Tissue samples were homogenized with a mini handheld homogenizer in lysis buffer (50 mM Tris, 150 mM NaCl, 5 mM EDTA, 1% triton X-100, 0.1% sodium deoxycholate) and centrifuged for 10 min at 5000*g*. The Pierce BCA Protein Assay Kit (Thermo Scientific cat. 23225) was used to determine the protein concentration of the supernatant of each sample.

The measurement of glucocerebrosidase activity in each homogenate sample was performed as previously described (Arrant et al., 2019). Each sample was diluted to 1 mg/mL in citrate phosphate buffer (0.1 M sodium citrate dihydrate, 0.2 M sodium phosphate monohydrate), such that 5 μg of protein (5 μL) was loaded in pentaplicate into a black, clear-bottom 96-well plate. Glucocerebrosidase assay buffer (citrate phosphate buffer with 1% bovine serum albumin, 0.25% triton X-100, 0.25% taurocholic acid, and 1.11 mM EDTA at pH 4.6) containing either 4-methylumbelliferone β-D-glucopyranoside (4-MUGluc, 1.11 mM) or 4-MUGluc and conduritol B epoxide (CBE, 0.2 mM) was added to each well at 45 μL such that three wells contained 4-MUGluc and two wells contained 4-MUGluc and CBE in a final volume of 50 μL. Each plate was analyzed with a set of blanks (no protein) to determine the background signal of 4-MUGluc, 4-MUGluc, and CBE. Reaction plates were incubated at 37℃ for 1 hr, after which 50 μL of stop solution (0.4 M glycine, pH 11) was added to each well. Standard dilutions of 4-MU, eight concentrations ranging from 0 – 7.5 μM in 100 μL of 50% assay buffer and 50% stop solution, were added to each plate in duplicate. Fluorescence intensity was recorded on a Synergy 2 plate reader (BioTek) at ex/em wavelengths of 360 nm/440 nm.

Specific glucocerebrosidase activity was determined for each sample by correcting for background fluorescence and the fluorescent signal produced in the presence of the glucocerebrosidase inhibitor, CBE. The corrected fluorescent signal of each sample was then compared to the standard curve of 4-MU to determine the nmol 4-MU produced per hour per mg protein loaded. Glucocerebrosidase activity was determined for the midbrain, hippocampus, striatum, and frontal cortex of 8 GBA1^+/+^ (6M:2F) and 8 GBA1^+/L444P^ mice (6M:2F)

### Immunoblotting

At three months of age, GBA1^+/+^ and GBA1^+/L444P^ mice were transcardially perfused with 0.9% NaCl saline. The brain was removed and dissected to isolate the midbrain, hippocampus, striatum, and cortex. The tissues were homogenized with a mini handheld homogenizer in homogenization buffer (1% Tx-100, protease inhibitor, and phosphatase inhibitor in TBS). Samples were centrifuged at 4°C for 30 min at 21,150*g* and supernatant was collected. The Pierce BCA Protein Assay Kit (Thermo Scientific cat. 23225) was used to determine the protein concentration of the supernatant of each sample.

Protein sample was loaded into each well of a 4-20% Mini-PROTEAN TGX Precast protein 15-well gels (Bio-Rad, cat#4561096). Protein was transferred using transfer buffer (20% MeOH in TBS) to a PVDF membrane overnight at 4 °C at 25 V. The next day, membranes were rinsed three times with TBS/1%Tween followed by one hour blocking using the Intercept® (TBS) Blocking Buffer and Diluent Kit (LI-COR, Cat#927-66003). After the membranes were rinsed three times using TBS-Tween, membranes were incubated in a primary solution from the diluent kit indicated above.

Primary antibodies used were as indicated: Vinculin (Bio-Rad Cat# MCA465GA, alternatively, Cat# MCA465S, RRID: AB_2214389), GCase (Sigma-Aldrich Cat# G4171, RRID: AB_1078958), Total α-syn (Abcam Cat# ab51252, RRID: AB_869971), phosphorylated α-syn (Abcam, Cat#ab51253, RRID: AB_869973), vGLUT1 (Synaptic Systems, Cat#135 304, RRID: AB_887878), Homer1 (Synaptic Systems Cat#160 006, RRID: AB_2631222), Tuj1 (Neuromics Cat# MO15013, RRID: AB_2737114), SNAP25 (Sigma-Aldrich Cat# S9684, RRID:AB_261576), VAMP2 (Synaptic Systems Cat# 104 211, RRID:AB_887811), Syntaxin-1a (Sigma-Aldrich Cat# SAB4502894, RRID:AB_10748204). After the membranes were incubated at 4 °C overnight, membranes were rinsed three times and then incubated in a secondary solution for 1 hour with the secondary diluent kit as indicated above. SDS was added to generate 0.01% SDS secondary antibody solution. The secondary antibodies were as indicated: IRDye® 800CW Goat anti-Rabbit IgG Secondary Antibody (LI-COR Biosciences Cat# 925-32211, RRID:AB_2651127), IRDye® 680RD Donkey anti-Guinea Pig IgG Secondary Antibody (LI-COR Biosciences Cat# 926-68077, RRID:AB_10956079), IRDye® 800CW Donkey anti-Chicken Secondary Antibody (LI-COR Biosciences Cat# 926-32218, RRID:AB_1850023), and AlexaFluor 680 anti-Mouse IgG Secondary Antibody. Bands were imaged and analyzed, by quantifying band intensity, using Image Studio Lite Ver 5.2. The band of the protein of interest was normalized to the loading control, vinculin. All immunoblots were run with 3-6 individual mice.

### Statistical Analysis

All data were graphed with GraphPad Prism 9 and analyzed with the ROUT test for outliers at Q = 10% then tested for normality using the Shapiro-Wilks test. Data that failed the test for normality were graphed as a scatter plot without a bar using a Mann-Whitney test. If data were normal, an independent t-test, two-way ANOVA, or Three-way ANOVA were used. All data are presented as mean ± SEM. A p<0.05 was considered statistically significant. A list of the statistical tests used for each graph can be found in **Table 1**.

**Table 1:**
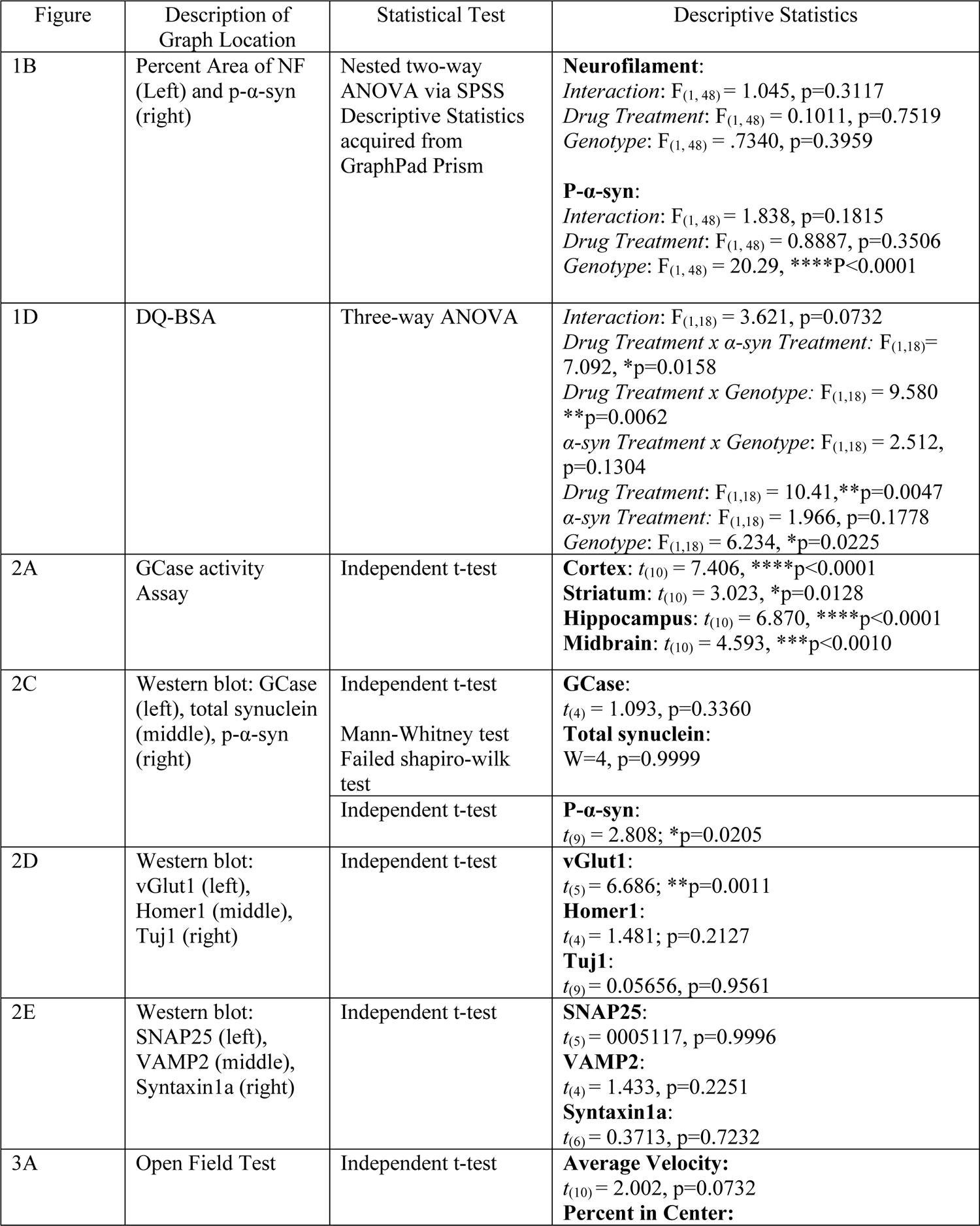
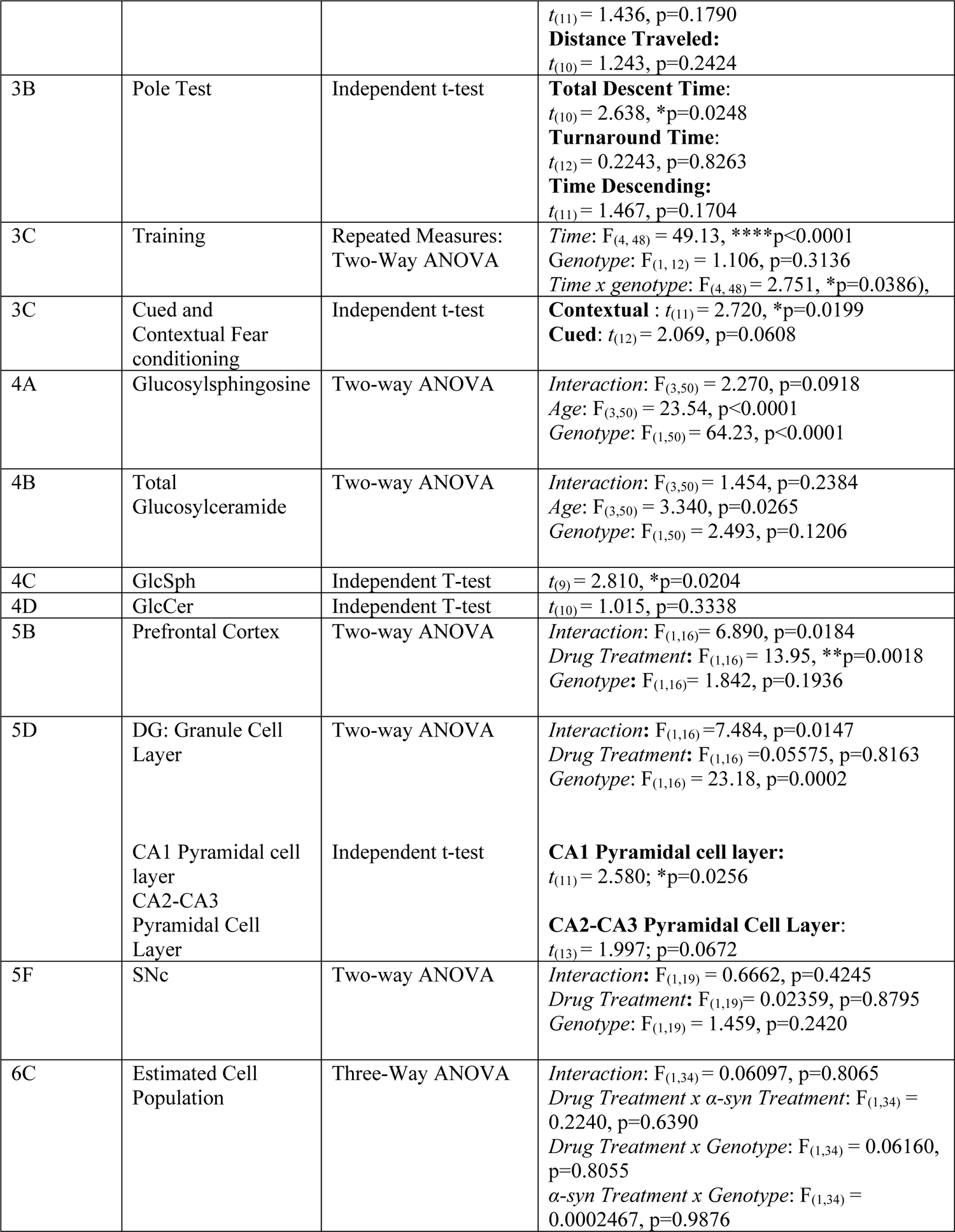
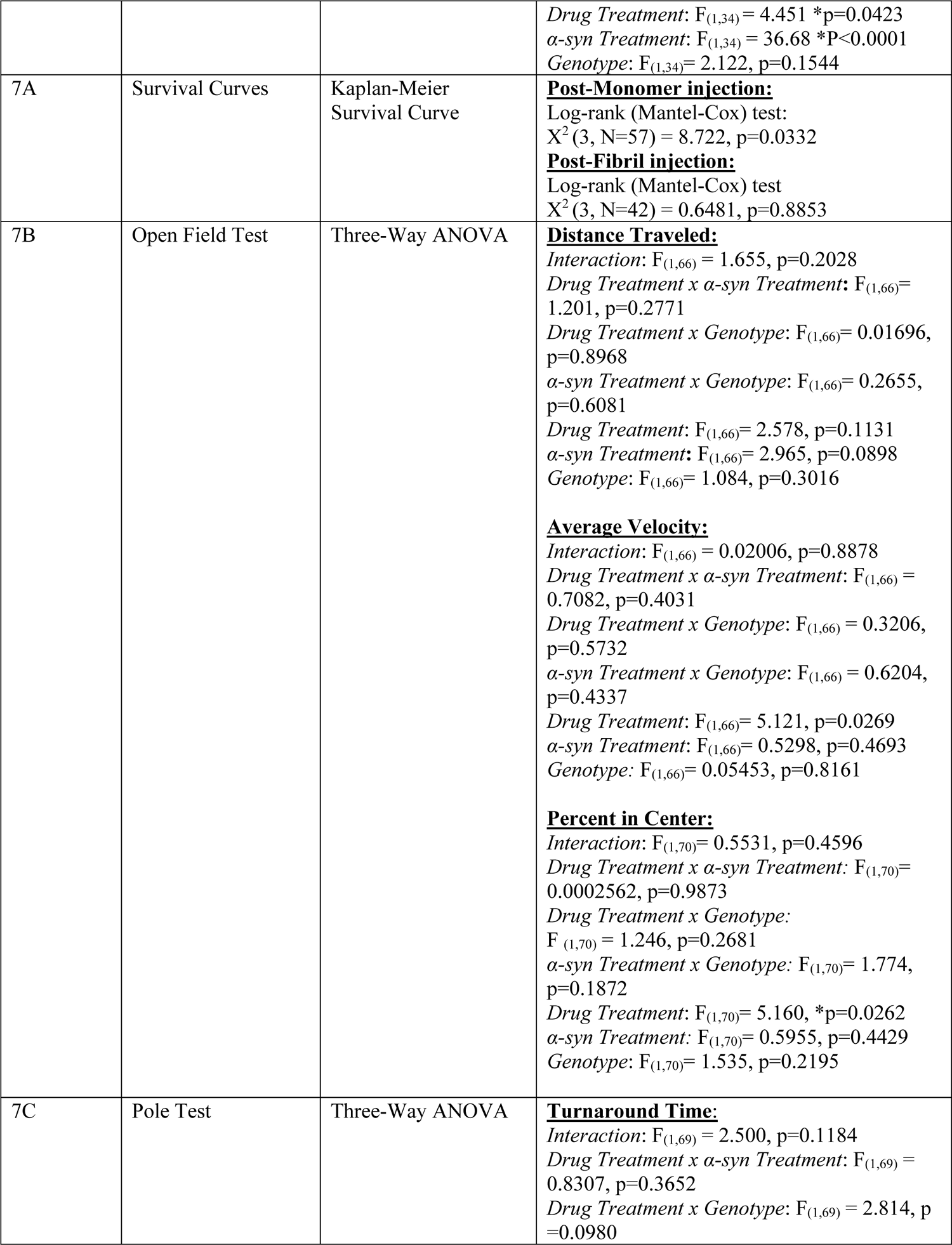
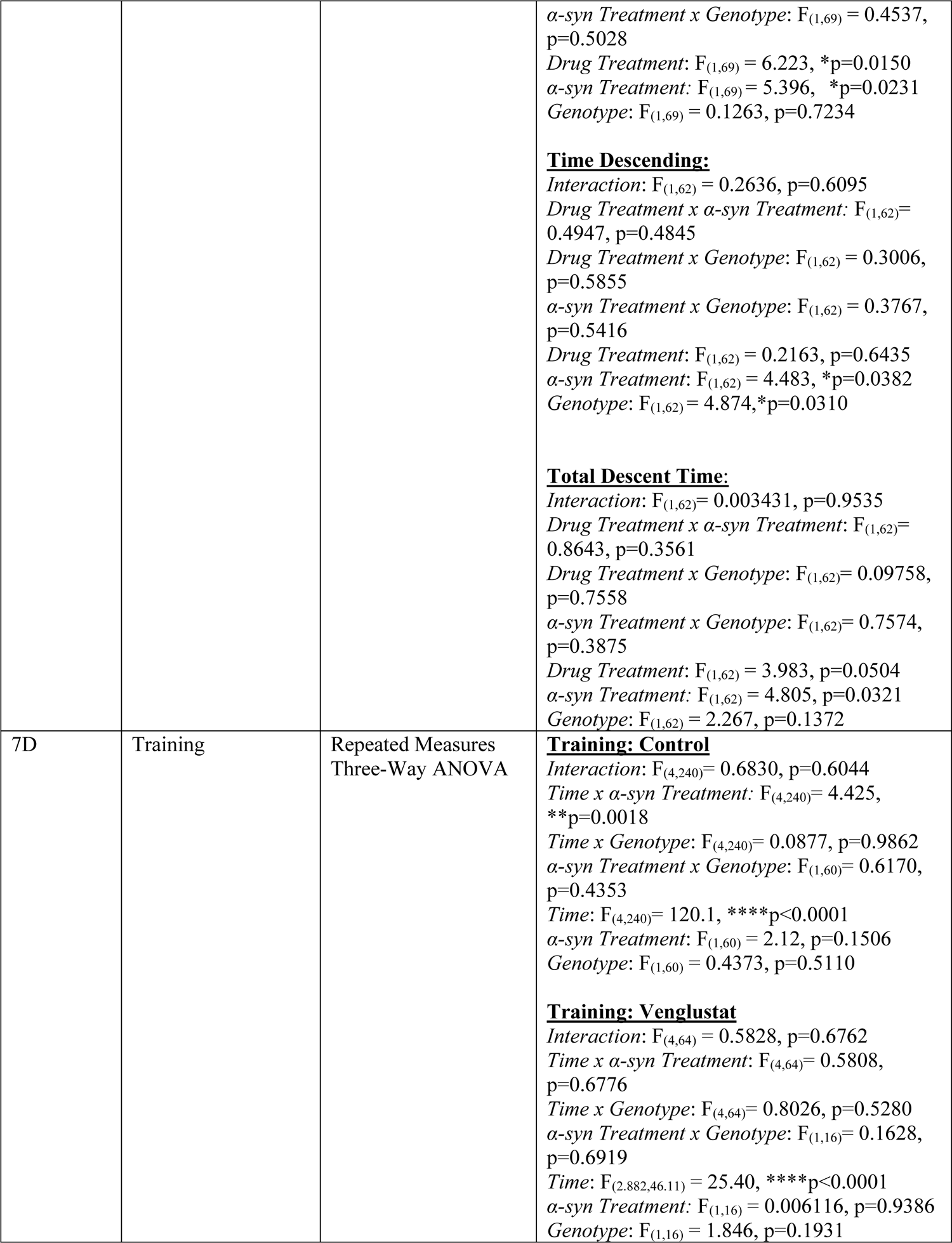
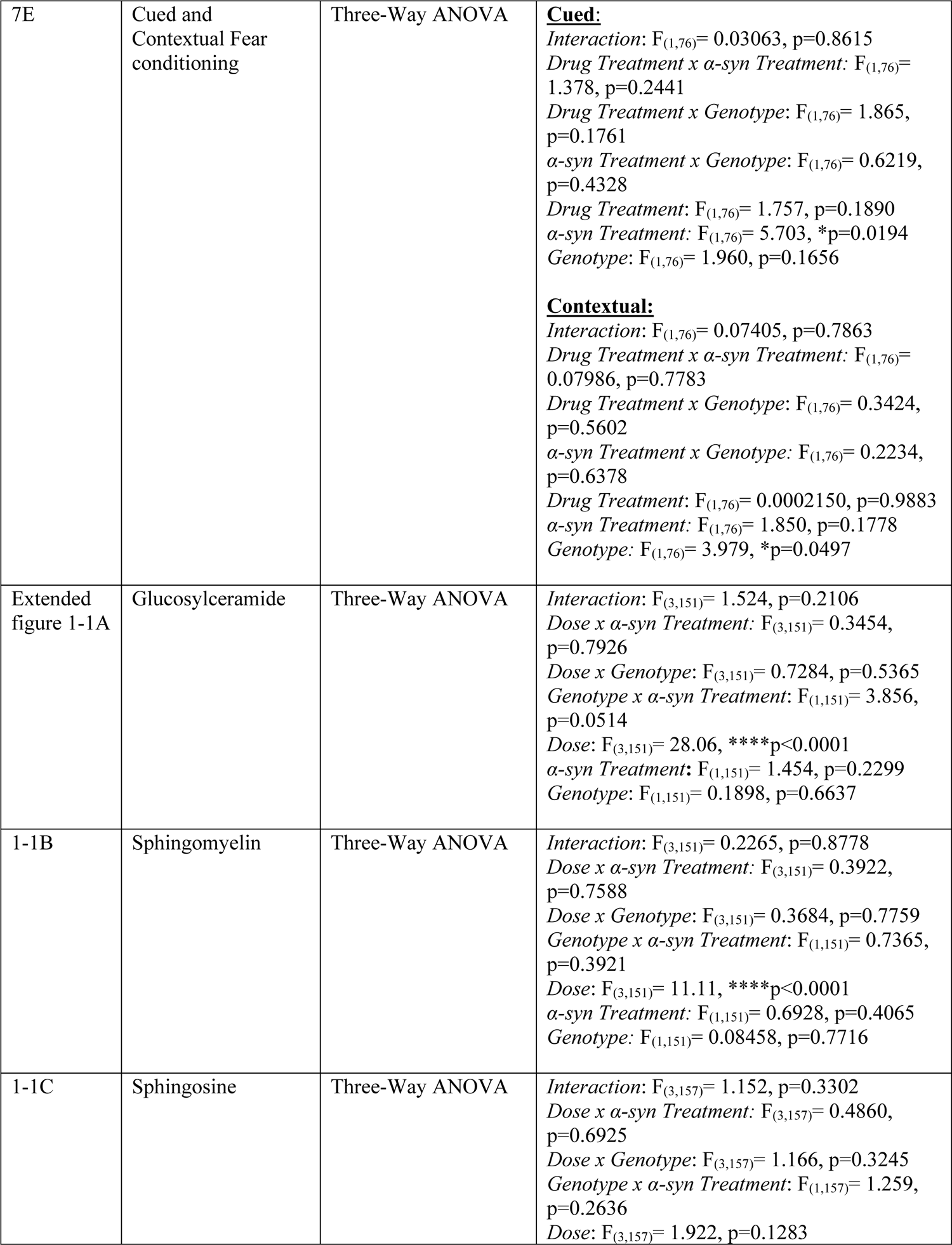
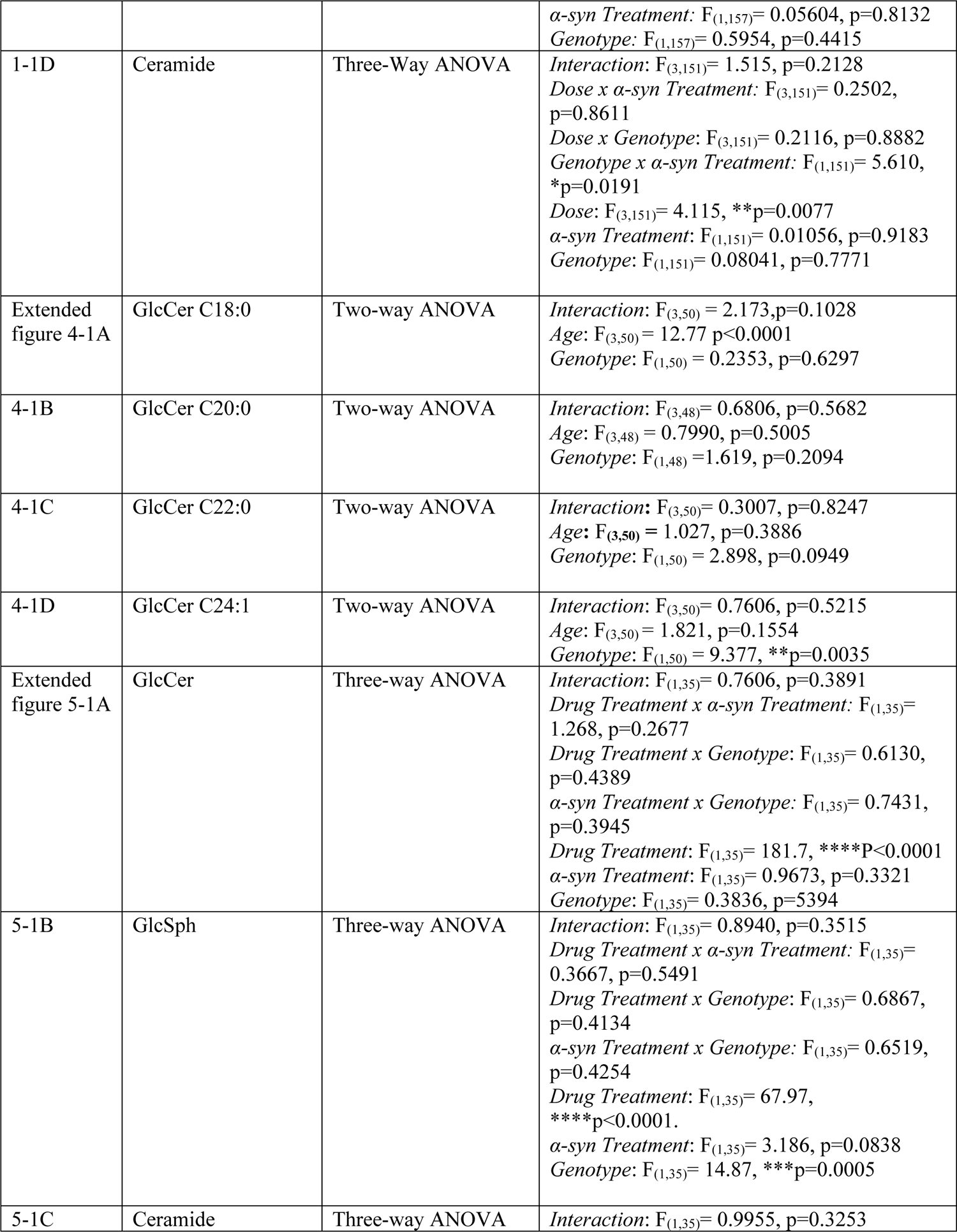
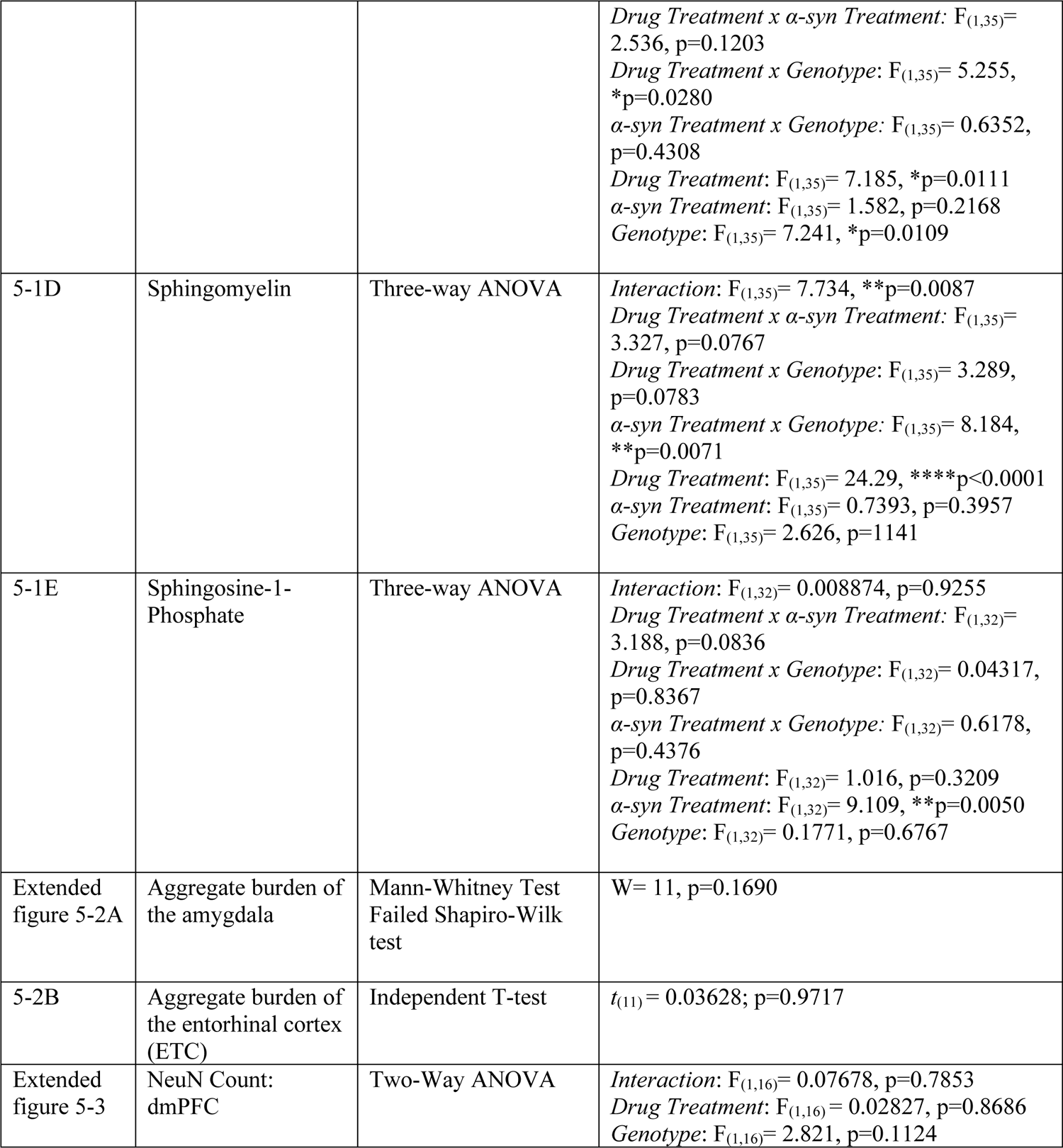
Statistical test for all graphs.

## RESULTS

### Fibril-induced α-syn inclusion formation in primary cultured hippocampal neurons from GBA1^+/+^ and GBA1^+/L444P^ mice

To study the effects of GBA1 L444P expression on α-syn inclusion formation, we first used primary hippocampal neurons from GBA1^+/+^ mice and GBA1^+/L444P^ knock-in mice exposed to sonicated α-syn fibrils on DIV7 and analyzed 14 days later (DIV21) (Volpicelli-Daley et al., 2011). Primary hippocampal neurons from mice were chosen as our model culture system because we achieve reliable, robust formation of α-syn inclusions in response to exposure to fibrils (Volpicelli-Daley et al., 2014; Froula et al., 2018; Brzozowski et al., 2021; Stoyka et al., 2021). Because the formation of α-syn inclusions depend on neuronal outgrowth (Murphy et al., 2000), we quantified axonal density using neurofilament-H (NFH; heavy polypeptide). There was no change in percent area occupied by NFH in neurons from GBA1^+/L444P^ mice compared to GBA1^+/+^ neurons with and without fibrils (**Figure 1a, 1b, left,** *DMSO:* p=0.9996 *Eliglustat*: p=0.2632). Fourteen days after exposure to fibrils, α-syn phosphorylated at Serine129 (p-α-syn), which is used as a marker of inclusion formation, was not significantly different between neurons from GBA1^+/L444P^ or GBA1^+/+^ mice (**Figure 1a, 1b, right,** *DMSO:* p=0.2149, *Eliglustat*: p<0.0001).

**Figure 1:**
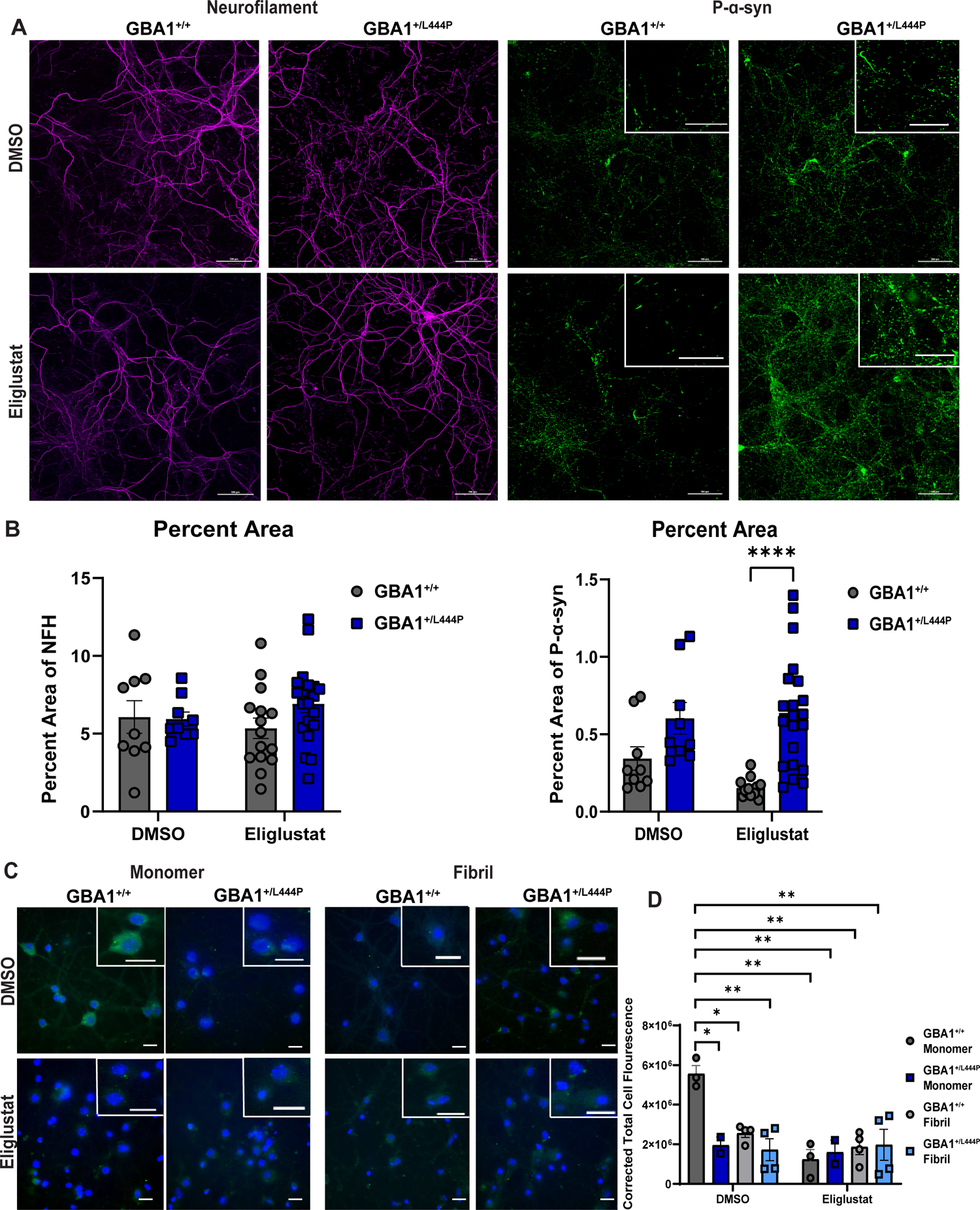
Primary hippocampal neurons from GBA1^+/L444P^ mice show reduced lysosome activity compared to those from GBA1^+/+^ mice and show increased α-syn inclusions 14 days after exposure to fibrils and eliglustat. A. Representative images of neurofilament (magenta, left) and p-α-syn (green, right) in hippocampal primary cultures. Top row of panels shows neurons treated with equivalent volumes of DMSO as Eliglustat (100 nM, bottom row of panels). Scale bar = 100µm, Zoom = 50µm. B. Quantification of the percent area of neurofilament (left; nested two-way ANOVA *Interaction*: *Time x Genotype*: F_(1, 49)_ = 1.365, p=0.2484, *Drug Treatment*: F_(1, 49)_ = 0.03061, p=0.8618, and *Genotype*: F_(1, 49)_ = 1.004, p=0.3212) and p-α-syn (right, nested two-way ANOVA; *Time x Genotype*: F_(1, 48)_ = 1.838, p=0.1815, *Drug Treatment*: F_(1, 48)_ = 0.8887, p=0.3506, and *Genotype*: F_(1, 48)_ = 20.29, ****p<0.0001, N=9 for GBA1^+/+^ and GBA1^+/L444P^ DMSO treated neuron groups, N=15 GBA1^+/+^ Eliglustat treated neurons, N=20 GBA1^+/L444P^ Eliglustat treated neurons). One GBA1^+/+^ DMSO and one GBA1^+/L444P^ DMSO outlier were removed for both NF and p-α-syn analysis based on Grubbs’ test. C. Representative images for DQ-BSA assay showing Hoechst (blue) and DQ Red BSA (visualized in green). Top row of panels shows neurons treated with equivalent volumes of DMSO as Eliglustat (100 nM, bottom row of panels) between GBA1^+/+^ and GBA1^+/L444P^ primary hippocampal neurons. D. Quantification of background corrected total cell fluorescence (Three-way ANOVA: *interaction*: F_(1,18)_ = 3.621, p=0.0732, *Drug Treatment x α-syn Treatment*: F_(1,18)_= 7.092, *p=0.0158, *Drug Treatment x Genotype*: F_(1,18)_ = 9.580, **P=0.0062, *α-syn Treatment x Genotype*: F_(1,18)_ = 2.512, P=0.1304, *Drug Treatment*: F_(1,18)_ = 10.41, **P=0.0047, *α-syn Treatment*: F_(1,18)_ = 1.966, P=0.1778, *Genotype*: F_(1,18)_ = 6.234, *P=0.0225; N=3 for GBA1^+/+^ Monomer DMSO and Eliglustat treated neuron groups, N=2 for GBA1^+/L444P^ Monomer DMSO and Eliglustat treated neuron groups, N=4 for GBA1^+/+^ and GBA1^+/L444P^ fibril DMSO and Eliglustat treated neurons groups). For all graphs, error bars represent SEM. Scale bar =100µm. *<0.05, **<0.01, and ****<0.0001.

Quantitation of sphingolipids by mass spectrometry in the primary neuronal cultures showed no differences in levels of glucosylceramide, total sphingomyelin, sphingosine, or ceramide in neurons exposed to fibrils compared to no treatment, or in neurons expressing GBA1^+/L444P^ compared to GBA1^+/+^ (**Extended figure 1-1a-d**). Glucosylsphingosine was not detectable in the primary neuronal cultures.

Because it has been proposed that mutant GBA1 increases α-syn aggregation by increasing levels of glucosylceramide (Mazzulli et al., 2011; Stojkovska et al., 2019), primary hippocampal neurons were treated with eliglustat which prevents the formation of glucosylceramide and glucosylsphingosine by inhibiting glucosylceramide synthase (GCS) (Bennett and Turcotte, 2015). Neurons were treated with 1nM, 10nM, and 100nM eliglustat (IC_50_ =20nM of intact MDCK cells, (Shayman, 2010)), and mass spectrometry quantitation of lipids showed that 100nM of eliglustat significantly reduced levels of total glucosylceramide, with no effect on total sphingomyelins, sphingosine, or ceramide (**Extended figure 1-1a-d**). Fourteen days of treatment with eliglustat did not alter axon density as seen with NF immunofluorescence (**Figure 1a, 1b, left**). However, fourteen days of eliglustat treatment significantly increased the abundance of fibril-induced α-syn inclusions in primary neurons from GBA1^+/L444P^ mice compared to neurons from GBA1^+/+^ mice **Figure 1a, 1b, right**).

### Lysosome function in primary cultured hippocampal neurons from GBA1^+/+^ mice or GBA1^+/L444P^ mice with and without fibrils

GCase activity plays a role in lysosome function (Boer et al., 2020). Thus, a DQ-BSA assay was used to determine the impact of GBA1 L444P on lysosome activity in neurons. According to this assay, active proteases in lysosomes cleave DQ-BSA, which then fluoresces (**Figure 1c, d**). Primary neurons from GBA1^+/L444P^ mice showed significantly reduced DQ-BSA fluorescence compared to neurons from GBA1^+/+^ mice. In addition, fourteen days of fibril exposure significantly reduced DQ-BSA fluorescence in neurons from GBA1^+/+^ mice compared to neurons from GBA1^+/+^ mice not exposed to fibrils. Fibril exposure did not further reduce DQ-BSA fluorescence in neurons from GBA1^+/L444P^ mice compared to GBA1^+/+^ mice. Eliglustat treated primary neurons showed significantly reduced DQ-BSA fluorescence compared to GBA1^+/+^ control neurons. Thus, while both expressions of GBA1 L444P and exposure to fibrils reduced lysosome activity, eliglustat treatment and reduction of glucosylceramide did not rescue these impairments.

### Expression of synaptic proteins and behavior in GBA1^+/L444P^ mice compared to GBA1^+/+^ mice

Because there are few studies characterizing baseline behavior and possible changes in the brains of GBA1^+/L444P^ mice, analyses of levels of neuronal protein expression and behavior in three-month-old GBA1^+/L444P^ mice compared to GBA1^+/+^ mice without fibrils or any stereotactic injections was performed. The activity of GCase in the cortex, striatum, hippocampus, and midbrain was analyzed; all regions showed reduced GCase activity in GBA1^+/L444P^ mice compared to GBA1^+/+^ mice. The reduction of GCase activity, as measured by the 4-MU assay, was 31.9% in the cortex, 21.6% in the striatum, 27.4% in the hippocampus, and 27.7% in the midbrain (**Figure 2a**). There was no change in the expression of levels of GCase protein (**Figure 2b, c**). In addition, there were no changes in levels of total synuclein, but rather a reduction in soluble p-α-syn between the GBA1^+/+^ and GBA1^+/L444P^ (**Figure 2b, c**). Hippocampal levels of synaptic and neuronal proteins were also analyzed. Levels of presynaptic vGLUT1, which transports glutamate into synaptic vesicles, were significantly reduced in the GBA1^+/L444P^ mice (**Figure 2b, d**) compared to GBA1^+/+^ mice. Levels of neuron-specific Tuj1 (beta-tubulin II) were unchanged between the two mouse genotypes (**Figure 2d**). There were no differences between GBA1^+/L444P^ mice and GBA1^+/+^ mice in levels of synaptic vesicle SNARE proteins, VAMP2, syntaxin1a, or SNAP-25. Overall, these data show a selective reduction in p-α-syn and vGLUT1 in GBA1^+/L444P^ without changes in other neuronal or synaptic proteins.

**Figure 2:**
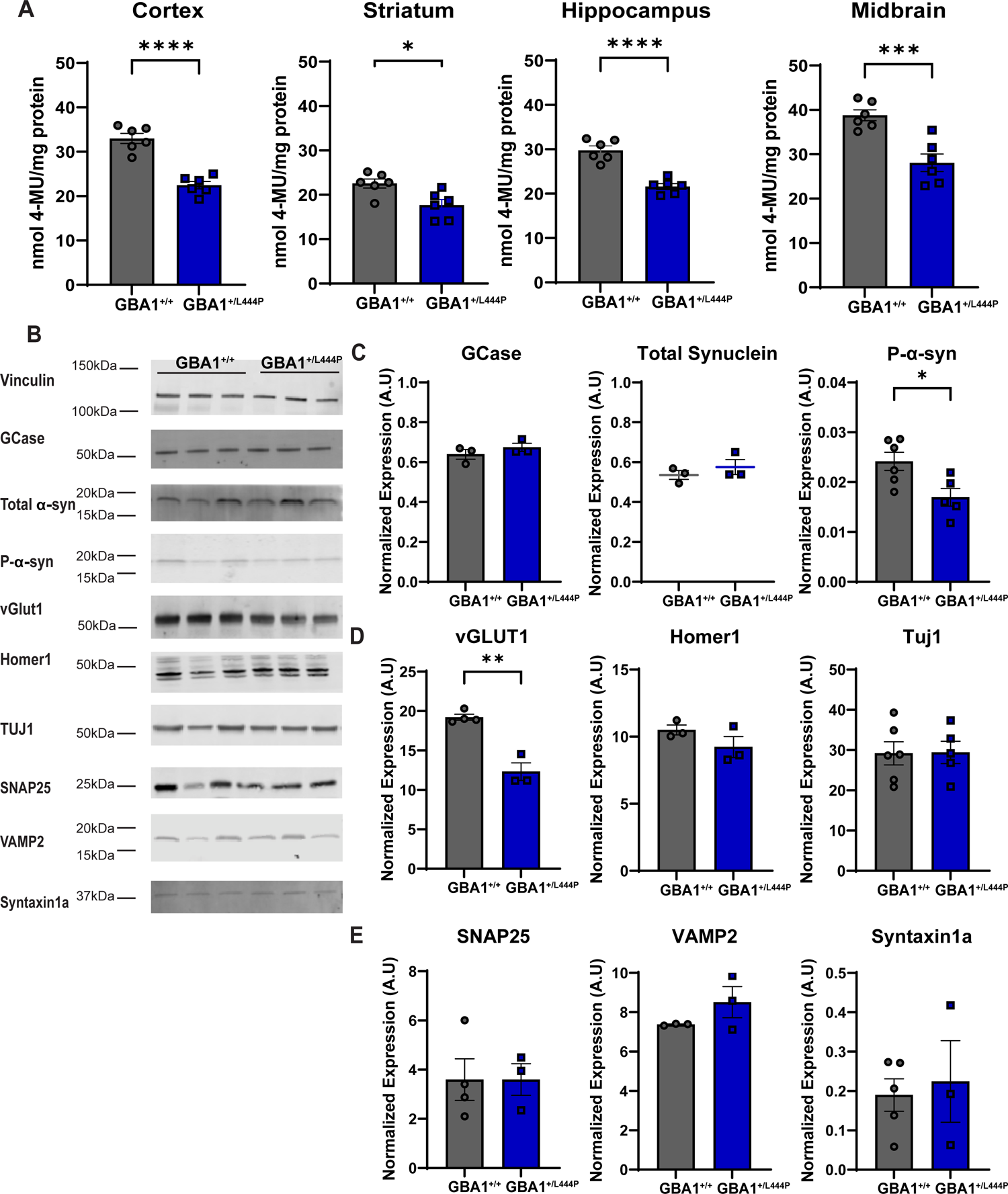
Brain homogenates from GBA1^+/L444P^ mice showed reduced GCase activity and selective reduction of vGLUT1 compared to GBA1^+/+^mice. A. GCase activity assay to evaluate enzymatic activity of GCase in two-month old GBA1^+/L444P^ mice in the cortex (independent t-test: *t*_(10)_ = 7.406, ****p<0.0001), striatum (*t*_(10)_ = 3.023, *p=0.0128), hippocampus (*t*_(10)_ = 6.870, ****p<0.0001), and the midbrain (*t*_(10)_ = 4.593; ***p<0.0010) expressed as nMol 4-MU/mg protein. N=6 for all groups. B. Representative immunoblot analyses of hippocampal brain lysates for GCase, endogenous total α-syn, p-α-syn, Homer1, TUJ1, SNAP25, VAMP2, and Syntaxin-1 expression. C. Quantitation of immunoblots; vinculin was used as the loading control to normalize expression. Independent t-test for GCase (*t*_(4)_ = 1.093, p=0.3360, N=3 for both groups, one GBA1^+/+^ outlier removed) and Mann-Whitney U test for endogenous total α-syn (W=4, p=0.9999,N=3 for both groups). An independent t-test was used for p-α-syn (*t*_(9)_ = 2.808; *p=0.0205, N= 6 GBA1^+/+^ and N=5 GBA1^+/L444P^). Analysis of vGLUT1 (Independent t-test: *t*_(5)_ = 6.686; **p=0.0011, N=4 GBA1^+/+^ and N=3 GBA1^+/L444P^, One GBA1^+/+^ outlier removed based on Grubbs’ test), Homer1 (*t*_(4)_ = 1.481; p=0.2127, N=3 both groups), and TUJ1 (*t*_(9)_ = 0.05656, p=0.9561, N=6 GBA1^+/+^ and N=5 GBA1^+/L444P^) expression. E. Analysis of SNAP25 (Independent t-test: *t*_(5)_ = 0005117, p=0.9996, N=4 GBA1^+/+^ and N=3 GBA1^+/L444P^), VAMP2 (*t*_(4)_ = 1.433, p=0.2251, N=3 in both groups, one GBA1^+/+^ outlier removed), and Syntaxin-1a (*t*_(6)_ = 0.3713, p=0.7232, N=5 GBA1^+/+^ and N=3 GBA1^+/L444P^) expression. All immunoblots were ran with 3-6 individual mice. For all graphs, error bars represent SEM. The Total Synuclein data failed the test for normality and graphed as a scatter plot without a bar using a Mann-Whitney test. *p<0.05, **p<0.01, ***p<0.001, and ****<0.0001.

Behavioral tests were also performed in the GBA1^+/L444P^ mice to determine performance in motor activity and associative learning. During the open field test, which evaluates general motor behavior, GBA1^+/L444P^ mice showed no statistical differences in the average velocity, total distance traveled, or the time spent in the center (**Figure 3a**). The pole test is used to test basal ganglia motor activity and slowness of movement reflected as increased time to descend pole, in models of PD (Hernandez et al., 2021). GBA1^+/L444P^ mice showed a faster total descent time compared to GBA1^+/+^ mice (**Figure 3b, left**), but there was no difference in turnaround time or the time descending the pole (**Figure 3b, middle and right**). Lastly, fear conditioning was performed to measure associative learning, previously shown to be impaired in mice with fibril-induced inclusions (Stoyka et al., 2020, 2021). During the training phase, both groups of mice showed increased freezing over time in response to a tone paired with a shock with approximately 73.6% and 73.8%% time freezing after foot shock for GBA1^+/+^ and GBA1^+/L444P^ mice, respectively (**Figure 3c, left**). The next day, memory and fear responses were tested by measuring the time spent freezing. In cued fear conditioning, mice were placed in a novel environment and exposed to the CS used for training. GBA1^+/L444P^ mice and GBA1^+/+^ mice exhibited no statistical difference in the percentage of time spent freezing (*t*_(12)_ = 2.069; p=0.0608) (**Figure 3c, middle**). In contextual fear conditioning, mice were placed in the same environment as the training without the presence of the tone. GBA1^+/L444P^ mice froze less than GBA1^+/+^ mice (**Figure 3c, right**). Despite variability between GBA1^+/L444P^ mice, no weight or percent time freezing differences between sex were observed (data not shown). Thus, the GBA1^+/L444P^ mice showed impairment in contextual fear memory.

**Figure 3:**
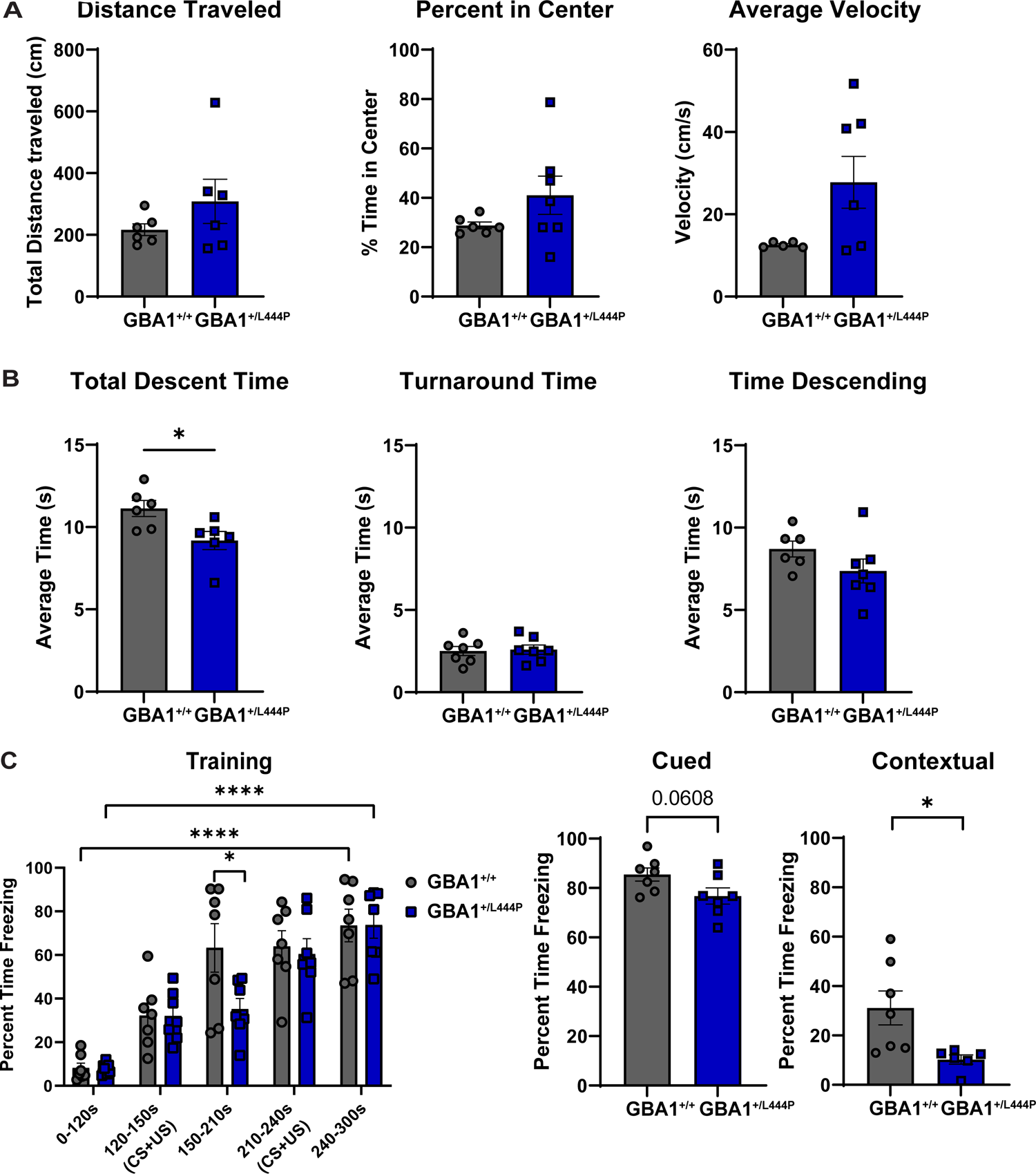
Behavioral analysis of three-month old GBA1^+/L444P^ mice compared to GBA1^+/+^ mice. **A**. Open field test analysis evaluating average velocity (independent t-test: 2 GBA1^+/+^ outliers removed; *t*_(10)_ = 2.002; p=0.0732, N=6 for both groups), percent in the center (one GBA1^+/+^ outlier removed; *t*_(11)_ = 1.436, p=0.1790, N=6 GBA1^+/+^ N=7 GBA1^+/L444P^), and distance traveled (one GBA1^+/+^ and GBA1^+/L444P^ outlier removed; *t*_(10)_ = 1.243, p=0.2424, N=5 GBA1^+/+^ N=7 GBA1^+/L444P^). **B**. Pole test evaluated total descent time (one GBA1^+/+^ and GBA1^+/L444P^ outlier removed: *t*_(10)_ = 2.638, *p=0.0248, N=6 for both groups), turnaround time, (*t*_(12)_ = 0.2243, p=0.8263, N=7 for both groups), and time descending (one GBA1^+/+^ outlier removed; *t*_(11)_ = 1.467; p=0.1704, N=6 GBA1^+/+^ N=7 GBA1^+/L444P^). **C**. Fear conditioning analysis evaluated percent time spent freezing in training (Repeated Measures Two-way ANOVA: *time*: F_(4, 48)_ = 49.13, ****p<0.0001, *genotype*: F_(1, 12)_ = 1.106, p=0.3136, *time x genotype*: F_(4, 48)_ = 2.751, *p=0.0386), cued fear conditioning (Independent t-test: *t*_(12)_ = 2.069, p=0.0608), and contextual fear conditioning (*t*_(11)_ = 2.720, *p=0.0199, one GBA1^+/L444P^ outlier removed). N=7 for both groups. *p<0.05, ***p<0.001, and ****<0.0001.

### Levels of glucosylceramides and glucosylsphingosine in GBA1^+/L444P^ mice and GBA1^+/+^ mice at different ages

Defects in sphingolipid metabolism due to reduced GCase activity are prevalent in Gaucher’s disease and have been implicated in PD (Mazzulli et al., 2011; Belmatoug et al., 2017; Taguchi et al., 2017). Thus, we evaluated the accumulation of GlcCer and GlcSph in the forebrain in 3-, 6-, 9-, and 12-months GBA1^+/+^ mice and GBA1^+/L444P^ mice. In GBA1^+/+^ mice, levels of GlcSph were significantly increased in 12-month-old mice compared to younger mice. GlcSph levels were significantly higher in the brains of GBA1^+/L444P^ mice compared to GBA1^+/+^ mice by 6 months of age. GlcSph levels increased with age in the GBA1^+/L444P^ mice as well (**Figure 4a**). In contrast, GlcCer levels remained unchanged between GBA1^+/+^ and GBA1^+/L444P^ mice (**Figure 4b**) To evaluate whether different acyl chain length structural variants of GlcCer were altered with age and the presence of the GBA1^+/L444P^ mutation, we evaluated four isoforms of GlcCer. GlcCer18:0 decreased with age in GBA1^+/+^ mice but reached a plateau at 9 months of age (**Extended Figure 4-1a**). However, there were no statistical differences between the GlcCer20:0, GlcCer22:0, and GlcCer24:1 isoform in either GBA1^+/+^ or GBA1^+/L444P^ mice (**Extended Figure 4-1b-d**).

**Figure 4:**
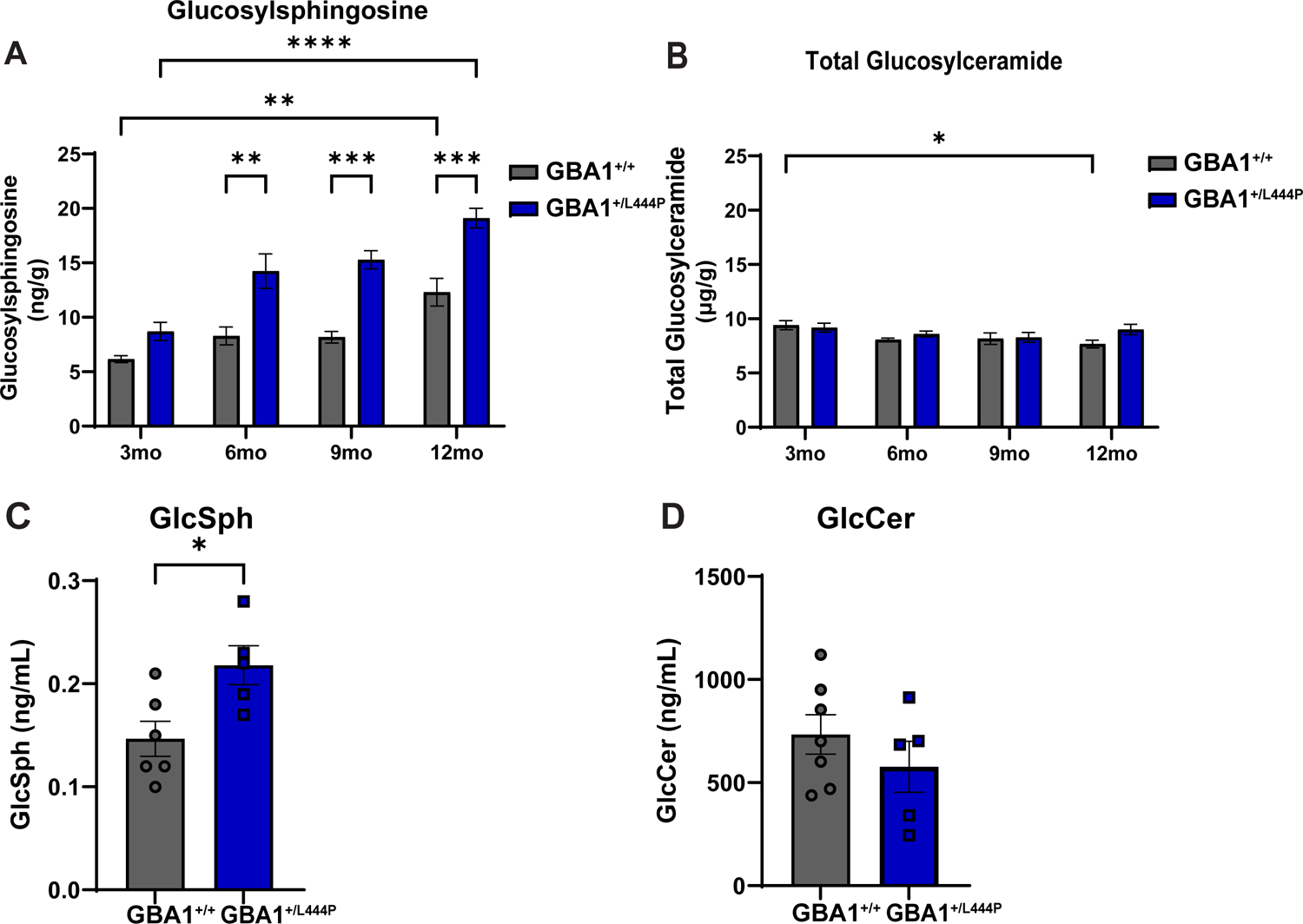
Lipid analysis in GBA1^+/+^ and GBA1^+/L444P^ mice. GBA1^+/+^ and GBA1^+/L444P^ mice forebrains underwent mass spectrometry for lipid quantification for **A.** Glucosylsphingosine (Two-way ANOVA; *Interaction*: F_(3,50)_ = 2.270, p=0.0918, *Age*: F_(3,50)_ = 23.54, ****p<0.0001, and *Genotype*: F_(1,50)_ = 64.23, ****p<0.0001) and **B.** total glucosylceramide (*Interaction*: F_(3,50)_ = 1.454, p=0.2384, *Age*: F_(3,50)_ = 3.340, *p=0.0265, *Genotype*: F_(1,50)_ = 2.493, p=0.1206) at 3mo, 6mo, 9mo and 12mo. **C.** Mice plasma was collected and underwent mass spectrometry for lipid quantification for **C**. GlcSph (Independent T-test, *t*_(9)_ = 2.810, *p=0.0204, N=6 for GBA1^+/+^, and N=5 GBA1^+/L444P^) and **D.** GlcCer (*t*_(10)_ = 1.015, p=0.3338, N=7 for GBA1^+/+^ and N=5 GBA1^+/L444P^). *p<0.05 and ****p<0.0001.

Recent studies have evaluated the effective use of lipid substrates in patient plasma as a potential biomarker of disease. Plasma GlcSph levels correlate with disease burden in Gaucher’s disease (Murugesan et al., 2016). Moreover, a recent study implemented the usefulness of GlcSph in GBA1-PD as a biomarker of disease (Surface et al., 2022). Thus, we evaluated the accumulation of lipids in the plasma of GBA1^+/+^ and GBA1^+/L444P^ mice at the time of sacrifice. Similar to findings in brain homogenates, GlcSph was increased in the plasma of GBA1^+/L444P^ mice compared to GBA1^+/+^, while GlcCer remained unchanged (**Figure 4c-d**).

### Fibril-induced α-syn inclusions in GBA1^+/L444P^ mice compared to GBA1^+/+^ mice

GBA1^+/+^ and GBA1^+/L444P^ mice were injected bilaterally in the dorsolateral striatum with fibrils or monomer at 2-3 months of age to induce α-syn inclusion formation. Starting one-week post injections until sacrifice, mice were fed chow containing venglustat or control chow to determine if inhibiting glucosylceramide synthase, and consequently reducing the synthesis of glycosphingolipids, reduces formation of α-syn inclusions. Ten months post injection, mice were perfused, and immunofluorescence was performed on fixed brain sections for p-α-syn positive inclusions. The dmPFC, hippocampus, and SNc were analyzed as previously described (Stoyka et al., 2020, 2021). Mice injected with monomer showed no p-α-syn inclusions, regardless of GBA1 L444P expression as previously shown in our publications (Froula et al., 2019; Stoyka et al., 2020). Abundant inclusions were seen in dmPFC, SNc, and the hippocampus in the fibril injected GBA1^+/+^ mice and GBA1^+/L444P^ mice. There were no significant differences in the pathological p-α-syn load in the dmPFC between genotypes in mice that received control chow (**Figure 5a, b**). In the hippocampus, GBA1^+/L444P^ mice had significantly more p-α-syn inclusions than GBA1^+/+^ mice in the control treatment group in the dentate gyrus and CA1 pyramidal cell layer **(Figure 5c, d)**. Finally, there were no significant differences or patterns in pathological load between GBA1 L444P expression and treatment conditions in the SNc (**Figure 5e, f**), basolateral amygdala, and entorhinal cortex (**Extended figure 5-1**).

**Figure 5:**
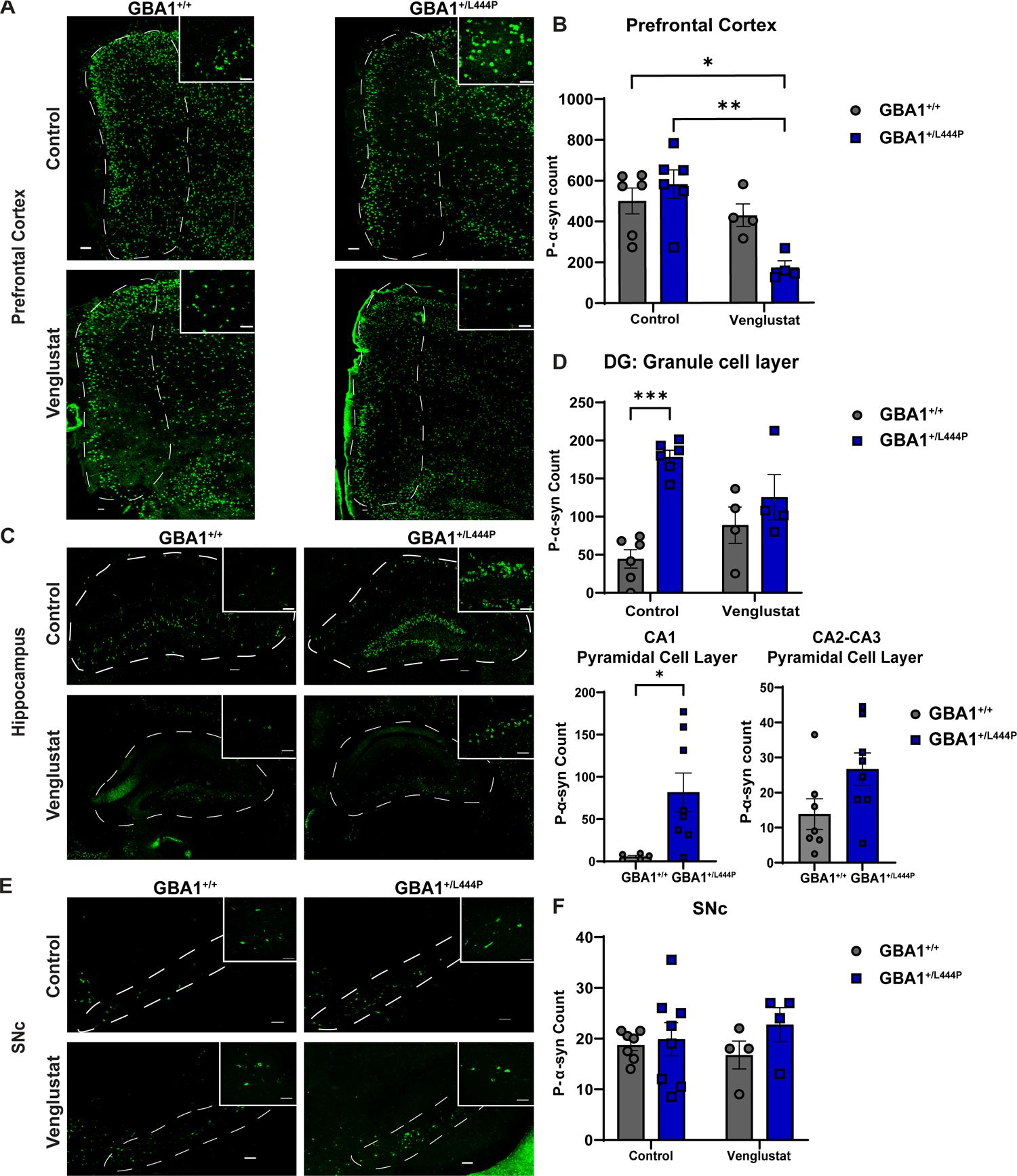
P-α-syn pathology burden is increased in the hippocampal formation. **A**. 10-months after bilateral striatal injections, immunofluorescence for p-α-syn was performed. Representative images of GBA1^+/+^ and GBA1^+/L444P^ fibril-injected mice fed control or venglustat chow of the dmPFC were captured using confocal microscopy. Scale bar =100um, zoomed photos scale bar = 50um. **B**. Quantification of p-α-syn in the dmPFC (N=6 for both control chow groups, N=4 for venglustat chow groups). Two-way ANOVA: *Interaction*: F_(1,16)_= 6.890, *p=0.0184, *Drug Treatment*: F_(1,16)_ = 13.95, **p=0.0018, *Genotype*: F_(1,16)_= 1.842, p=0.1936. Error bars represent SEM. **C**. Representative images of GBA1^+/+^ and GBA1^+/L444P^ fibril-injected mice fed control or venglustat chow of the hippocampus were captured using confocal microscopy. Scale bar =100um, zoomed photos scale bar = 50um. **D**. Quantification of p-α-syn in the granule cell layer of the dentate gyrus area of the hippocampus (N=6 for both control chow groups, N=4 for venglustat chow groups, Two outliers removed (One GBA1^+/+^ and one GBA1^+/L444P^). *Interaction*: F_(1,16)_ =7.484, *p=0.0147, *Drug Treatment*: F_(1,16)_ =0.05575, p=0.8163. *Genotype*: F_(1,16)_ = 23.18, ***p=0.0002. Quantification of CA1-CA3 pyramidal cell layer with control chow (N=7 GBA1^+/+^ and N=8 GBA1^+/L444P^: T-test: CA1 Pyramidal cell layer: *t*_(11)_ = 2.580; *p=0.0256, CA2-CA3 Pyramidal Cell Layer: *t*_(13)_ = 1.997; p=0.0672) Error bars represent SEM. **E**. Representative images of GBA1^+/+^ and GBA1^+/L444P^ fibril-injected mice fed control or venglustat chow of the SNc were captured using confocal microscopy. Scale bar =100um, zoomed photos scale bar = 50um. **F**. Quantification of p-α-syn in the SNc (N=7 for GBA1^+/+^ and N=8 for GBA1^+/L444P^ control chow groups, N=4 for GBA1^+/+^ and GBA1^+/L444P^ venglustat chow groups). Two-way ANOVA: *Interaction*: F_(1,19)_ = 0.6662, p=0.4245, *Drug Treatment*: F_(1,19)_= 0.02359, p=0.8795, *Genotype*: F_(1,19)_ = 1.459, p=0.2420. *p<0.05, **p<0.01, and ***p<0.001.

To determine if inhibiting GCS prevented α-syn inclusion formation, mice were provided with chow with venglustat or control chow. To ensure the biological activity of venglustat treatment, plasma was acquired from the mice groups at the time of sacrifice, and mass spectrometry was performed to measure GlcCer concentrations. Compared to control mice, GlcCer was significantly reduced in all venglustat treated mice (**Extended figure 5-2a**). Mice administered venglustat showed significantly reduced plasma levels of GlcCer, confirming the efficacy of the compound. However, GBA1^+/+^ mice did not show a difference in abundance of α-syn inclusions between mice treated with or without venglustat (**Figure 5**). GBA1^+/L444P^ mice treated with venglustat did show significantly fewer inclusions in the mPFC than both GBA1^+/+^ mice and GBA1^+/L444P^ mice that were not treated with venglustat (**Figure 5a, b**). This difference was not due to cell death as there were no changes in the quantity of NeuN^+^ cells between any groups (**Extended Figure 5-3**). Treatment with venglustat did not reduce the abundance of α-syn inclusions in the hippocampus (**Figure 5c, d**) or in the SNc (**Figure 5e, f**).

### Neuroinflammation in GBA1^+/L444P^ mice compared to GBA1^+/+^ mice

Expression of the GBA1 L444P mutation causes systemic inflammation in addition to enhanced neuroinflammation clinically and in this mouse model (Mizukami et al., 2002; Rocha et al., 2015; Mus et al., 2019; Brunialti et al., 2021; Williams et al., 2021). Thus, ten months post-monomer or fibril injection, hippocampal brain sections were stained for MHCII and IgG to investigate inflammation. Despite the increased accumulation of pathologic α-syn in fibril-injected GBA1^+/L444P^ mice, the granule cell layer of the dentate gyrus showed no differences between IgG and MHCII abundance across all groups (**Extended figure 5-4**).

### Fibril-induced loss of TH-positive dopaminergic neurons in the SNc in GBA1^+/+^ mice and GBA1^+/L444P^ mice

Fibril-induced formation of α-syn inclusions leads to the death of dopamine neurons in the SNc (Luk et al., 2012; Froula et al., 2019; Stoyka et al., 2021). To test whether GBA1^+/L444P^ mice show enhanced dopaminergic loss, immunohistochemistry and unbiased stereology were performed using antibodies to TH to quantify SNc dopamine neurons. Both GBA1^+/+^ mice and GBA1^+/L444P^ mice injected with fibrils had an approximately 50% reduction of dopaminergic neurons in the SNc compared to GBA^+/+^ mice injected with α-syn monomer (**Figure 6a and 6c**) as previously shown (Froula et al., 2019; Stoyka et al., 2020, 2021). Similarly, GBA1^+/L444P^ fibril-injected mice treated with venglustat showed a 50% decrease in TH^+^ neurons compared to GBA1^+/+^ monomer injected mice fed venglustat. However, no differences between GBA1 L444P expression, nor between venglustat and control chow, were observed. Thus, venglustat did not protect SNc neurons from toxicity caused by fibril-induced α-syn inclusions (**Figure 6b and 6c)**.

**Figure 6:**
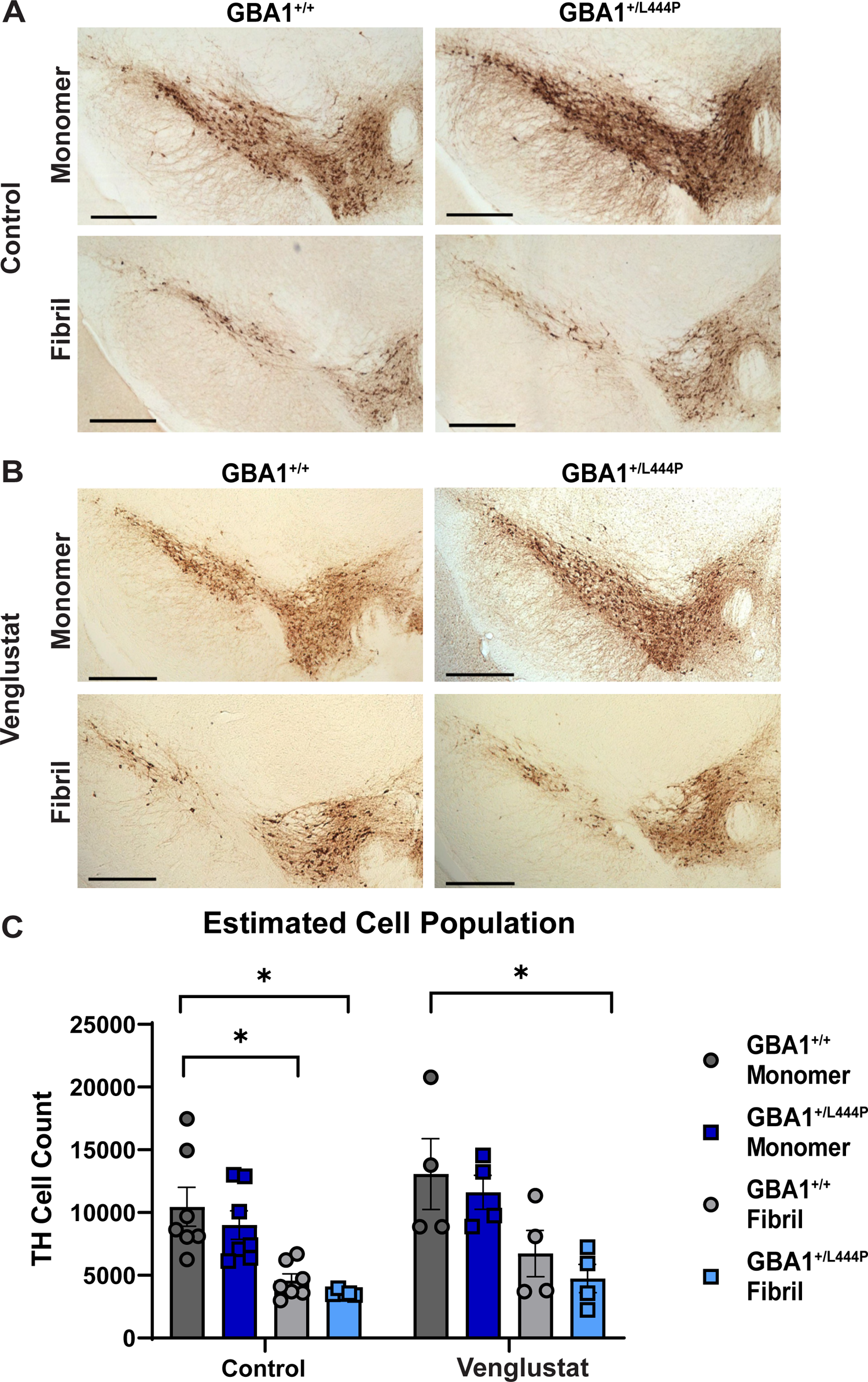
Quantification of Dopaminergic neuron cell death. Immunohistochemistry for TH was performed inn GBA1^+/+^ and GBA1^+/L444P^ mice at 10-months post injection, and the SNc was analyzed for dopaminergic cell death using unbiased stereology. **A.** Representative images of monomer and fibril injected GBA1^+/+^ and GBA1^+/L444P^ mice fed control chow (N=7 for all groups). **B**. Representative images of monomer and fibril injected GBA1^+/+^ and GBA1^+/L444P^ mice fed venglustat chow (N=4 for all groups). **C**. Unbiased stereology was used to estimate TH^+^ Cell count (Three-way ANOVA: *Interaction*: F_(1,34)_ = 0.06097, p=0.8065, *Drug Treatment x α-syn Treatment*: F_(1,34)_ = 0.2240, p=0.6390, *Drug Treatment x Genotype*: F_(1,34)_ = 0.06160, p=0.8055, *α-syn Treatment x Genotype*: F_(1,34)_ = 0.0002467, p=0.9876, *Drug Treatment*: F_(1,34)_ = 4.451 *p=0.0423, *α-syn Treatment*: F_(1,34)_ = 36.68 ****P<0.0001, *Genotype*: F_(1,34)_= 2.122, p=0.1544). Error bars represent SEM. Scale bar =200um. *p<0.05.

### Behavioral analyses in fibril injected GBA1^+/L444P^ mice compared to GBA1^+/+^ mice, with and without venglustat

Our lab previously showed that fibril-induced α-syn inclusions cause motor and nonmotor phenotypes (Froula et al., 2019; Stoyka et al., 2020). We evaluated how motor and nonmotor phenotypes are impacted in GBA1^+/L444P^ mice injected with fibrils and the effect of venglustat treatment. A Kaplan-Meier survival curve showed that GBA1^+/L444P^ monomer mice provided with venglustat chow died statistically significantly more often than the GBA1^+/+^ or GBA1^+/L444P^ monomer mice or the GBA1^+/+^ monomer-injected mice fed venglustat chow (**Figure 7a, left**). However, when comparing fibril-injected mice, there were no statistical differences between any groups (**Figure 7a, right**).

**Figure 7:**
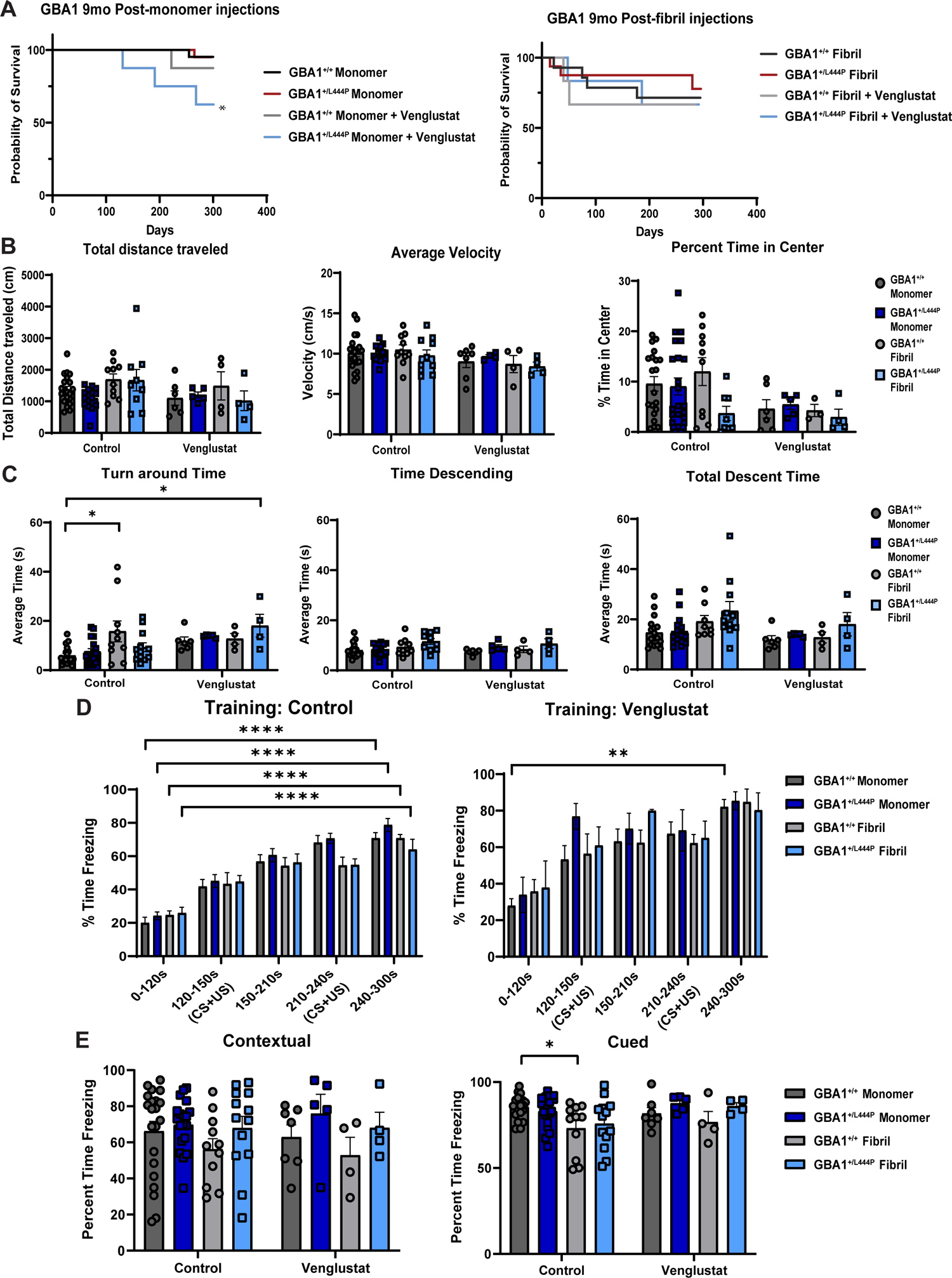
Behavioral analyses of monomer and fibril injected mice fed venglustat or control chow. GBA1^+/+^ and GBA1^+/L444P^ mice underwent behavioral test to evaluate motor and cognitive function seven months post-fibril or monomer injection and venglustat or control treatment. **A.** Quantification of mouse survival using a Kaplan-Meier survival curve (Log-rank (Mantel-Cox) test: post-monomer injection: Х^2^ (3, N=57) = 8.722, *p=0.0332, post-fibril injection: Х^2^ (3, N=42) = 0.6481, p=0.8853). **B**. Open field test analysis evaluating the total distance traveled (Three-way ANOVA: *Interaction*: F_(1,66)_ = 1.655, p=0.2028, *Drug Treatment x α-syn Treatment*: F_(1,66)_= 1.201, p=0.2771, *Drug Treatment x Genotype*: F_(1,66)_= 0.01696, p=0.8968, *α-syn Treatment x Genotype*: F_(1,66)_= 0.2655, p=0.6081, *Drug Treatment*: F_(1,66)_= 2.578, p=0.1131, *α-syn Treatment*: F_(1,66)_= 2.965, p=0.0898, *Genotype*: F_(1,66)_= 1.084, p=0.3016, Control: Monomer-injected GBA1^+/+^ N= 20, Monomer-injected GBA1^+/L444P^ N=16, Fibril-injected GBA1^+/+^ N= 10 (one outlier removed), Fibril-injected GBA1^+/L444P^ N=9. Venglustat: Monomer-injected GBA1^+/+^ N= 6, Monomer-injected GBA1^+/L444P^ N=5, and Fibril-injected GBA1^+/+^ and GBA1^+/L444P^ N= 4), average velocity (*Interaction*: F_(1,66)_ = 0.02006, p=0.8878, *Drug Treatment x α-syn Treatment*: F_(1,66)_ = 0.7082, p=0.4031, *Drug Treatment x Genotype*: F_(1,66)_ = 0.3206, p=0.5732, *α-syn Treatment x Genotype*: F_(1,66)_ = 0.6204, p=0.4337, *Drug Treatment*: F_(1,66)_= 5.121, *p=0.0269, *α-syn Treatment*: F_(1,66)_= 0.5298, p=0.4693, *Genotype*: F_(1,66)_= 0.05453, p=0.8161, Control: Monomer-injected GBA1^+/+^ N= 20, Monomer-injected GBA1^+/L444P^ N=16 (two outlier removed), Fibril-injected GBA1^+/+^ and GBA1^+/L444P^ N=10. Venglustat: Monomer-injected GBA1^+/+^ N= 7, Monomer-injected GBA1^+/L444P^ N=4, Fibril-injected GBA1^+/+^ and GBA1^+/L444P^ N= 4), and the percent time in the center (Interaction: F_(1,70)_= 0.5531, p=0.4596, *Drug Treatment x α-syn Treatment*: F_(1,70)_= 0.0002562, p=0.9873, *Drug Treatment x Genotype*: F _(1,70)_ = 1.246, p=0.2681, *α-syn Treatment x Genotype*: F_(1,70)_= 1.774, p=0.1872, *Drug Treatment*: F_(1,70)_= 5.160, *p=0.0262, *α-syn Treatment*: F_(1,70)_= 0.5955, p=0.4429, *Genotype*: F_(1,70)_= 1.535, p=0.2195, Control: Monomer-injected GBA1^+/+^ N= 20, Monomer-injected GBA1^+/L444P^ N=22 (one outlier removed), Fibril-injected GBA1^+/+^ N= 10 (one outlier removed), Fibril-injected GBA1^+/L444P^ N=8 (two outliers removed). Venglustat: Monomer-injected GBA1^+/+^ N= 6, Monomer-injected GBA1^+/L444P^ N=5, Fibril-injected GBA1^+/+^ N= 3 (one outlier removed), Fibril-injected GBA1^+/L444P^ N=4). **C.** Mice underwent pole test to evaluate motor deficits by analyzing the time descending, (Three way ANOVA: *Interaction*: F_(1,62)_ = 0.2636, p=0.6095, *Drug Treatment x α-syn Treatment*: F_(1,62)_= 0.4947, p=0.4845, *Drug Treatment x Genotype*: F_(1,62)_ = 0.3006, p=0.5855, *α-syn Treatment x Genotype*: F_(1,62)_ = 0.3767, p=0.5416, *Drug Treatment*: F_(1,62)_ = 0.2163, p=0.6435, *α-syn Treatment*: F_(1,62)_ = 4.483, *p=0.0382, *Genotype*: F_(1,62)_ = 4.874, *p=0.0310, Control: Monomer-injected GBA1^+/+^ N= 17, Monomer-injected GBA1^+/L444P^ N=15, Fibril-injected GBA1^+/+^ N= 9, Fibril-injected GBA1^+/L444P^ N=10. Venglustat:, Monomer-injected GBA1^+/+^ N= 6, Monomer-injected GBA1^+/L444P^ N=5, Fibril-injected GBA1^+/+^ and GBA1^+/L444P^ N= 4), turnaround time (*Interaction*: F_(1,69)_ = 2.500, p=0.1184, *Drug Treatment x α-syn Treatment*: F_(1,69)_ = 0.8307, p=0.3652, *Drug Treatment x Genotype*: F_(1,69)_ = 2.814, p=0.0980, *α-syn Treatment x Genotype*: F_(1,69)_ = 0.4537, p=0.5028, *Drug Treatment*: F_(1,69)_ = 6.223, *p=0.0150, *α-syn Treatment*: F_(1,69)_ = 5.396, *p=0.0231, *Genotype*: F_(1,69)_ = 0.1263, p=0.7234; Control: Monomer-injected GBA1^+/+^ N= 18, Monomer-injected GBA1^+/L444P^ N=18, Fibril-injected GBA1^+/+^ N= 10, Fibril-injected GBA1^+/L444P^ N=12. Venglustat: Monomer-injected GBA1^+/+^ N= 6, Monomer-injected GBA1^+/L444P^ N=5, Fibril-injected GBA1^+/+^ and GBA1^+/L444P^ N= 4), and total descent time (*Interaction*: F_(1,62)_= 0.003431, p=0.9535, *Drug Treatment x α-syn Treatment*: F_(1,62)_= 0.8643, p=0.3561, *Drug Treatment x Genotype*: F_(1,62)_= 0.09758, p=0.7558, *α-syn Treatment x Genotype*: F_(1,62)_= 0.7574, p=0.3875, *Drug Treatment*: F_(1,62)_ = 3.983, p=0.0504, *α-syn Treatment*: F_(1,62)_ = 4.805, *p=0.0321, *Genotype*: F_(1,62)_ = 2.267, p=0.1372; Control: Monomer-injected GBA1^+/+^ N= 17, Monomer-injected GBA1^+/L444P^ N=15, Fibril-injected GBA1^+/+^ N= 8, Fibril-injected GBA1^+/L444P^ N=11. Venglustat: Monomer-injected GBA1^+/+^ N= 6, Monomer-injected GBA1^+/L444P^ N=5, Fibril-injected GBA1^+/+^ and GBA1^+/L444P^ N= 4). **D**. Quantification of the training phase of fear conditioning for monomer or fibril-injected GBA1^+/+^ or GBA1^+/L444P^ mice fed control chow (Repeated measures three-way ANOVA: *Interaction*: F_(4,240)_= 0.6830, p=0.6044, *Time x α-syn Treatment:* F_(4,240)_= 4.425, **p=0.0018, *Time x Genotype*: F_(4,240)_= 0.0877, p=0.9862, *α-syn Treatment x Genotype*: F_(1,60)_= 0.6170, p=0.4353, *Time*: F_(4,240)_= 120.1, ****p<0.0001, *α-syn Treatment*: F_(1,60)_ = 2.12, p=0.1506, *Genotype*: F_(1,60)_ = 0.4373, p=0.5110, GBA1^+/+^ Monomer N=21, GBA1^+/L444P^ Monomer N=19, GBA1^+/+^ fibril N= 10, GBA1^+/L444P^ fibril =13) and venglustat chow (*Interaction*: F_(4,64)_ = 0.5828, p=0.6762, *Time x α-syn Treatment*: F_(4,64)_= 0.5808, p=0.6776, *Time x Genotype*: F_(4,64)_= 0.8026, p=0.5280, *α-syn Treatment x Genotype*: F_(1,16)_= 0.1628, p=0.6919, *Time*: F_(2.882,46.11)_ = 25.40, ****p<0.0001, *α-syn Treatment:* F_(1,16)_ = 0.006116, p=0.9386, *Genotype*: F_(1,16)_ = 1.846, p=0.1931, GBA1^+/+^ Monomer N=7, GBA1^+/L444P^ Monomer N= 6, GBA1^+/+^ fibril N= 4, GBA1^+/L444P^ fibril =4). **E**. Quantification of cued (Three-way ANOVA: *Interaction*: F_(1,76)_= 0.03063, p=0.8615, *Drug Treatment x α-syn Treatment:* F_(1,76)_= 1.378, p=0.2441, *Drug Treatment x Genotype*: F_(1,76)_= 1.865, p=0.1761, *α-syn Treatment x Genotype*: F_(1,76)_= 0.6219, p=0.4328, *Drug Treatment*: F_(1,76)_= 1.757, p=0.1890, *α-syn Treatment:* F_(1,76)_= 5.703, *p=0.0194, *Genotype*: F_(1,76)_= 1.960, p=0.1656) Control: GBA1^+/+^ Monomer N=22, GBA1^+/L444P^ Monomer N=19, GBA1^+/+^ fibril N=9 (One outlier removed), GBA1^+/L444P^ fibril N=14, Venglustat: GBA1^+/+^ Monomer N=6, GBA1^+/L444P^ Monomer N=4, GBA1^+/+^ fibril N=4, GBA1^+/L444P^ fibril N=4), and contextual fear conditioning for all mice groups. Contextual Fear conditioning analysis was analyzed by evaluating the first 60 seconds (*Interaction*: F_(1,76)_= 0.07405, p=0.7863, *Drug Treatment x α-syn Treatment:* F_(1,76)_= 0.07986, p=0.7783, *Drug Treatment x Genotype*: F_(1,76)_= 0.3424, p=0.5602, *α-syn Treatment x Genotype:* F_(1,76)_= 0.2234, p=0.6378, *Drug Treatment*: F_(1,76)_= 0.0002150, p=0.9883, *α-syn Treatment:* F_(1,76)_= 1.850, p=0.1778, *Genotype:* F_(1,76)_= 3.979, *p=0.0497), Control: GBA1^+/+^ Monomer N=18, GBA1^+/L444P^ Monomer N=19, GBA1^+/+^ fibril N=9 (One outlier removed), GBA1^+/L444P^ fibril N=13, Venglustat: GBA1^+/+^ Monomer N=7, GBA1^+/L444P^ Monomer N=5, GBA1^+/+^ fibril N=4, GBA1^+/L444P^ fibril N=4). Error bars represent SEM. *p<0.05, **p<0.01, and ****<0.0001.

To evaluate motor behavior, an open field test was performed and showed no statistical differences in any groups for total distance traveled, or average velocities. In addition, there were no differences among groups with respect to percent time in the center. Although the venglustat treated group showed less time in the center (Drug Treatment: F_(1,70)_= 5.160, *p=0.0262), the Tukey’s post-hoc test detected no statistical differences between groups (**Figure 7b**). For the pole test, GBA1^+/+^ mice injected with fibrils and GBA1^+/L444P^ mice injected with fibrils and fed venglustat chow spent increased time turning around on the pole compared to GBA1^+/+^ mice injected with monomer, indicative of reduced motor function consistent with previous literature (**Figure 7c, left**) (Froula et al., 2019).

Lastly, to test for associative learning, fear conditioning was used as described in figure 3. During the training phase, all mice fed control chow displayed an increase in the percent of time spent freezing ranging from 63%-78%. (**Figure 7d, left)**. However, in venglustat treated mice, only GBA1^+/+^ mice injected with monomeric α-syn displayed an increase in freezing of approximately 82% (**Figure 7d, right).** No other differences between control and venglustat fed mice were observed (data not shown).

The next day, mice were tested for memory by going through cued and contextual fear conditioning as described in figure 2. For contextual fear conditioning, mice freezing behavior was examined during the first minute. However, all mice groups displayed equal percent time freezing from 52% to 75% across venglustat treated groups and 56% to 69% in control-treated mice groups (**Figure 7e left**). In cued fear conditioning, only GBA1^+/+^ fibril-injected mice fed control chow reduced by 12.43% in freezing behavior compared to GBA1^+/+^ Monomer injected mice fed control chow. There were no differences among any other groups (**Figure 7e, left**).

## DISCUSSION

Heterozygosity for the severe GBA1 L444P mutation increases the risk of PD dementia (Cilia et al., 2016). However, the mechanisms by which mutant GBA1 could increase neuronal susceptibility are relatively unclear. In this work, we show that levels of GlcSph, but not levels of GlcCer, increased in GBA1^+/L444P^ mice, supporting rising evidence that GlcSph is a highly relevant biomarker for GBA1-PD (Taguchi et al., 2017). Interestingly, aged GBA1^+/+^ mice also showed increased levels of GlcSph. Since age may be the primary risk factor for development of PD, these data indicate a potential role for GlcSph in the development of PD. In addition, neurons from GBA1^+/L444P^ mice showed reduced lysosome function, and reduced expression of vGLUT1, without changes in other synaptic proteins. Thus, heterozygous expression of GBA1 L444P alone, without Lewy pathology, may increase susceptibility of neurons by impacting lysosome and synapse biology. Our findings show, similar to previous studies, that heterozygous expression of GBA1 L444P increases pathologic α-syn, but only in the hippocampus of fibril-injected GBA1^+/L444P^ mice compared to fibril injected GBA1^+/+^ mice. Loss of dopaminergic neurons in the SNc was equivalent in GBA1^+/+^ and GBA1^+/L444P^ mice with fibril-induced α-syn inclusions, suggesting that GBA1 L444P mutations may selectively affect neurons in limbic brain areas. Pathologic inclusion formation and loss of SNc dopamine neurons were not rescued with inhibition of glucosylceramide synthase, supporting that GlcCer may not be the most relevant endpoint to monitor for GBA1-PD based therapeutics.

Heterozygosity of the GBA1 L444P mutation is moderately common and categorized as a severe mutation in GBA1-PD (Cilia et al., 2016). We demonstrate that the GBA1^+/L444P^ mutation alone leads to deficits also present in GBA1-PD. As such, we see reduced GCase activity in multiple brain regions including the cortex, striatum, hippocampus, and midbrain. This reduction of GCase activity may contribute to multiple outcomes also established. For instance, by three months of age, GBA1^+/L444P^ mice show reduced vGLUT1 in hippocampal brain lysates. Given how vGLUT1 is predominantly expressed in cortical and hippocampal regions (Wojcik et al., 2004), this reduction of vGLUT1 may underlie contextual associative memory deficits seen at three months of age. The contribution of GlcSph to pathologic outcomes is not fully understood. However, by 6-months of age, GBA^+/L444P^ mice show an increase in GlcSph in the forebrain of mice. Collectively, these data suggest that the mutation alone leads to impairments that are prevalent in GBA1-PD.

Given our findings that GCase activity is reduced in the hippocampus, cortex, and midbrain, we would expect an increase in aggregate burden in these regions as indicated in previous literature (Cullen et al., 2011; Mazzulli et al., 2011; Sardi et al., 2011; Migdalska-richards et al., 2020; Henderson et al., 2021). However, enhanced pathologic α-syn is only observed in the hippocampus, specifically in the granule cell layer of the dentate gyrus and the pyramidal cell layer of the CA1 region. Interestingly, under physiological conditions GCase is highly expressed in the hippocampus and entorhinal cortex (Dopeso-Reyes et al., 2018). Hippocampal neurons from GBA1^+/L444P^ mice show reduced lysosomal function.

Hippocampal primary cultures are enriched with excitatory neurons which show reduced expression of protein involved in degradative pathways (Fu et al., 2019). Thus, hippocampal neurons may be particularly susceptible to reduced GCase activity relative to other neuronal subtypes. Further studies examining GCase metabolic pathways in specific brain areas and neuronal populations could reveal how mutant GBA1 increases neuronal susceptibility in PD. However, we cannot rule out that using the fibril model, hippocampal pathology develops at later points than in other areas, and perhaps, the reason changes are only observed in the hippocampus is due to other regions reaching a plateau the in aggregate count by ten months post-injection.

Inhibition of GCS did not prevent α-syn inclusion formation in the hippocampus of fibril-injected mice despite showing previous activity in other central nervous system model systems at reducing disease burden (Marshall et al., 2016). Previous studies have shown that sphingolipids, predominantly GlcCer and GlcSph, accumulate coinciding with α-syn inclusion formation and reduced GCase activity (Mazzulli et al., 2011; Taguchi et al., 2017). However, GlcSph, but not GlcCer, accumulates in GBA1^+/L444P^ mice (Sardi et al., 2011; Taguchi et al., 2017). Moreover, when examining the various chain lengths for GlcCer, GlcCer18:0 was the most common isoform in the mice. A recent study showed that longer carbon lipid chains (>C22) lead to more pathologic outcomes (Fredriksen et al., 2021), and thus GlcCer18:0 in the mouse brain may not be impacted in this PD model. Overall, accumulating evidence points to aberrations in metabolism of GlcSph, and not GlcCer, as playing a pathologic role. This stresses the importance of evaluating other approaches to prevent the accumulation of GlcSph specifically. Future experiments could evaluate the inhibition of ASAH11, an inhibitor of GlcSph synthesis, or other mechanisms linked to GlcSph (Taguchi et al., 2017).

Multiple studies have demonstrated that inhibition of GCS prevents α-syn aggregate burden, GlcCer accumulation, and alleviates cognitive deficits (Sardi et al., 2017; Cosden et al., 2021; Viel et al., 2021). While our findings suggest that GCS inhibition alleviates aggregate burden in the mPFC, in general, our results fail to reproduce a robust effect of GCS inhibition on synucleinopathy *in vitro* or *in vivo*. One chief difference between the previous studies and our study is the distinction between the GBA1 L444P mutation and the GBA1 D409V mutation (Johnson et al., 2021). Interestingly, the heterozygous L444P mutation reduced GCase activity by 21.6-31.9% depending on the brain region evaluated. In contrast, the homozygous D409V mutation had a significantly greater reduction of GCase activity (Polinski et al., 2021). While the reduction of GCase activity may not be associated with PD risk or severity, it may contribute to the efficacy of GCS inhibition through other mechanisms (Omer et al., 2022). In addition, in GBA1^+/L444P^ cultures, increases in glucosylsphingosine were not detected unlike in GBA1^D409V/D409V^ cultures (Cosden et al., 2021). Suppressing GlcCer in the absence of elevated GlcSph may therefore not be sufficient to mitigate pathology. Moreover, there are structural and physiological differences between hippocampal and cortical neurons which may account for the variation in GCS inhibitor function.

However, further investigation is needed. There are also other important differences between methodologies between these studies. *In vitro*, even slight variations in pH could account for the GCS compound performing differently between studies. Moreover, there could be differences in the generation of fibrils between labs. Changes in the structure and size of fibrils result in different phenotypes and α-syn aggregation (Froula et al., 2019). Differences between animal models, such as transgenic expression of α-syn compared to fibril-induced corruption of endogenously expressed α-syn, and heterozygosity versus homozygosity of the mutation, may contribute to the variations seen between phenotypes observed (Sardi et al., 2017; Cosden et al., 2021; Viel et al., 2021). To further elaborate, the background strain of a mouse model is relevant in studying cognitive deficits. As such, C57BL/6 mice perform more efficiently than a 129/Sv mouse in the Morris water maze (Hall and Roberson, 2012). However, generating a mixed background of these two can produce a cognitive phenotype not seen in the pure 129/Sv mice (Wolfer et al., 1997). Additionally, sex difference is an important quantifier as female, but not male, heterozygous GBA1 D409V mice displayed a change in odor detection (Johnson et al., 2021). Collectively, therefore, we must note that there are multiple differences in experimental design which may give rise to these differential pathologic outcomes.

The enhanced pathologic burden of α-syn in the hippocampus, along with reduced GCase activity, may be a mechanistic explanation for the observed reduction of associative learning and memory function, especially related to clinically GBA1-PD cognitive function (Alcalay et al., 2010, 2012; Cilia et al., 2016; Fagan and Pihlstrøm, 2017). In summary, this study sheds light on the possible mechanisms by which mutant GBA1 may increase the risk for cognitive decline by influencing pathologic α-syn formation in the hippocampus. Further, our study highlights the potential importance of GlcSph, as a relevant biomarker for future therapeutics. Because GlcSph is also increased in the brains of aged mice, GlcSph targeted therapeutic approaches may be effective for both GBA1-PD and idiopathic PD. We also show, in mouse models, that levels of GlcSph in the plasma parallel those in the brains, suggesting this lipid holds value as a potential biomarker for PD. Thus, this study provides a pathway for understanding how GlcSph can impact brain function in aging and neurodegenerative disease as well as in the context of evaluating future therapeutic approaches to PD.

## Author contributions

CMC, MV, SMS, JNM and LVD designed research; CMC, MV, DR, R.AC.C, JF, SB, SC, MK, LY, B.-L.Y., NGH, performed research; CMC, MV, DR, SMS, JNM, and LVD wrote paper

The authors declare no competing financial interest.

## Acknowledgments

The authors wish to thank Valentina Krendelchtchikova for her help with purifying recombinant α-synuclein, Nicholas Boyle for help with protocols, and Dr. Karen Gamble and Dr. Richard Kennedy for advice on statistics. We also thank Dr. Joseph Mazzulli for the suggestion to characterize the GBA1 L444P mice without fibrils or injections. Lastly, we would like to thank the National Institutes of Health (NIH) National Institute of Neurological Disorders and Stroke (NINDS) Grant R01 NS102257 (to L.A.V.-D.), the NIH NINDS Grant R56 NS117465 (to L.A.V.-D.), the NIH NINDS Grant T32 NS095775 (to C.L.M.-C.), and the Parkinson’s Association of Alabama for funding this research.

