## Extended data for "Neuronopathic GBA1 L444P mutation accelerates Glucosylsphingosine levels and formation of hippocampal alpha-synuclein inclusions"

### Extended Figure 1-1

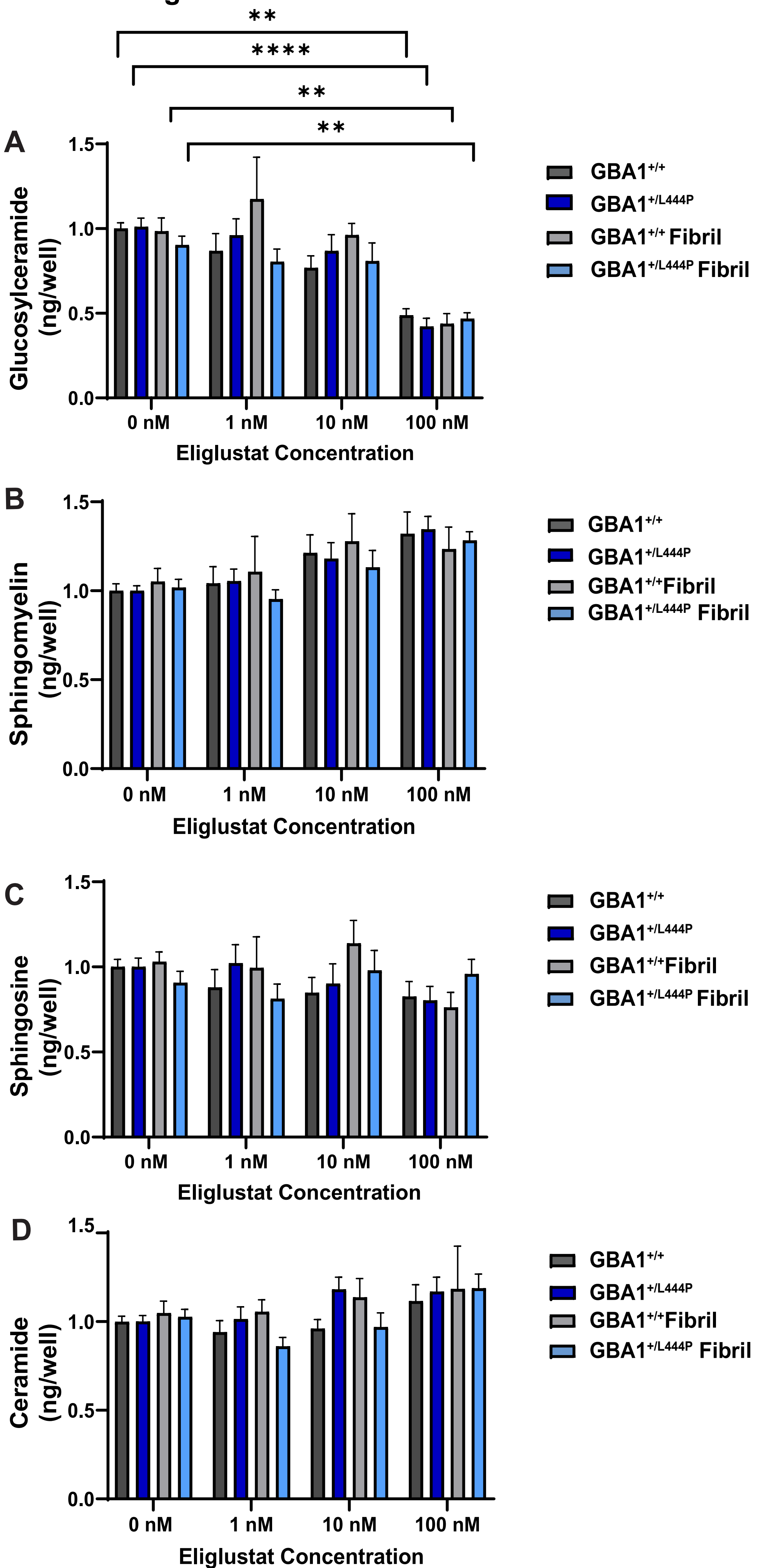

**Extended figure 1-1: Determination of Eliglustat Concentration based on lipid quantification**. GBA1$^{+/+}$ and GBA1$^{+/L444P}$ Primary cortical neurons were treated at DIV14 with varying concentrations of eliglustat (0, 1, 10, and 100 nM). Neurons were quantified by mass spectrometry for **A.** GlcCer (Three-way ANOVA; *Interaction*: $F_{(3,151)}$= 1.524, p=0.2106, *Dose x α-syn Treatment*: $F_{(3,151)}$= 0.3454, p=0.7926, *Dose x Genotype*: $F_{(3,151)}$= 0.7284, p=0.5365, *Genotype x α-syn Treatment*: $F_{(1,151)}$= 3.856, p=0.0514, *Dose*: $F_{(3,151)}$= 28.06, ****p<0.0001, *α-syn Treatment*: $F_{(1,151)}$= 1.454, p=0.2299, *Genotype*: $F_{(1,151)}$= 0.1898, p=0.6637) and **B.** sphingomyelin (*Interaction*: $F_{(3,151)}$= 0.2265, p=0.8778, *Dose x α-syn Treatment*: $F_{(3,151)}$= 0.3922, p=0.7588, *Dose x Genotype*: $F_{(3,151)}$= 0.3684, p=0.7759, *Genotype x α-syn Treatment*: $F_{(1,151)}$= 0.7365, p=0.3921, *Dose*: $F_{(3,151)}$= 11.11, ****p<0.0001, *α-syn Treatment*: $F_{(1,151)}$= 0.6928, p=0.4065, *Genotype*: $F_{(1,151)}$= 0.08458, p=0.7716,) **C**. sphingosine (*Interaction*: $F_{(3,157)}$= 1.152, p=0.3302, *Dose x α-syn Treatment*: $F_{(3,157)}$= 0.4860, p=0.6925, *Dose x Genotype*: $F_{(3,157)}$= 1.166, p=0.3245, *Genotype x α-syn Treatment*: $F_{(1,157)}$= 1.259, p=0.2636, *Dose*: $F_{(3,157)}$= 1.922, p=0.1283, *α-syn Treatment*: $F_{(1,157)}$= 0.05604, p=0.8132, *Genotype*: $F_{(1,157)}$= 0.5954, p=0.4415) and **D.** Ceramide (*Interaction*: $F_{(3,151)}$= 1.515, p=0.2128, *Dose x α-syn Treatment*: $F_{(3,151)}$= 0.2502, p=0.8611, *Dose x Genotype*: $F_{(3,151)}$= 0.2116, p=0.8882, Genotype x α-syn Treatment: $F_{(1,151)}$= 5.610, *p=0.0191, *Dose*: $F_{(3,151)}$= 4.115, **p=0.0077, *α-syn Treatment*: $F_{(1,151)}$= 0.01056, p=0.9183, *Genotype*: $F_{(1,151)}$= 0.08041, p=0.7771). Error bars represent SEM. For GlcCer, Sphingomyelin, and Ceramide in control treated neurons, GBA1$^{+/+}$ N=18, GBA1$^{+/L444P}$ N=24, fibril treated GBA1$^{+/+}$ N=18, and fibril treated GBA1$^{+/L444P}$ neurons N=24. To analyze sphingosine for control-treated neurons, GBA1$^{+/+}$ fibril-treated neurons N=24. For all lipid quantitation for eliglustat treated neurons (1nM, 10nM, and 100nM), GBA1$^{+/+}$ N=6, GBA1$^{+/L444P}$ N=8, fibril treated GBA1$^{+/+}$ N=6, and fibril treated GBA1$^{+/L444P}$ N=8, apart from fibril treated GBA1+/+ at 100nM where N=5. All sample sizes depict the number of individual mice. *p<0.05, **p<0.01, and ****<0.0001.

### Extended Figure 4-1

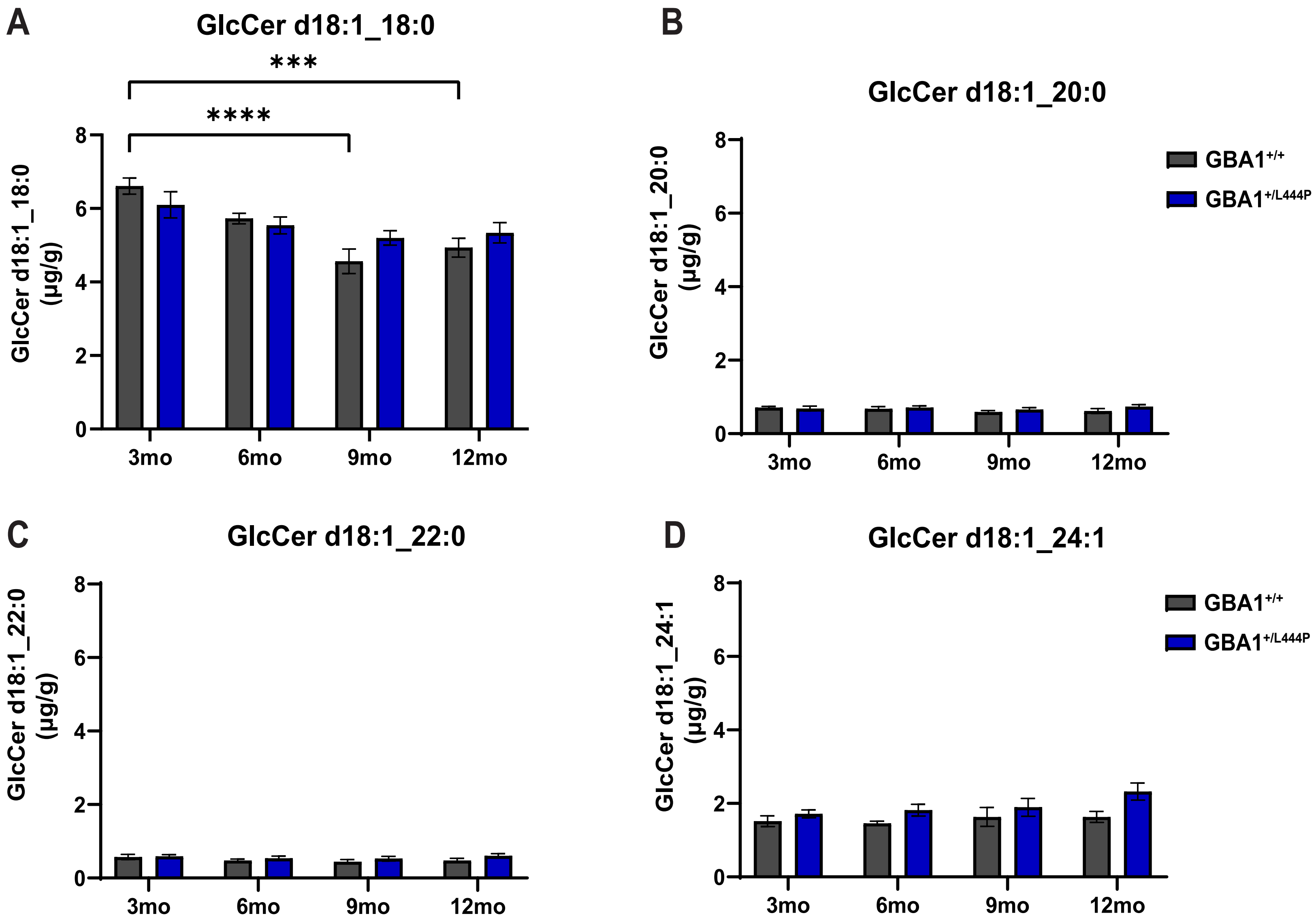

**Extended Data 4-1: Evaluation of GlcCer isoform quantification.** Four isoforms of GlcCer **A**. GlcCer C18:0 (Two-way ANOVA; *Interaction*: $F_{(3,50)} = 2.173$, p=0.1028, *Age*: $F_{(3,50)} = 12.77$, ****p<0.0001, and *Genotype*: $F_{(1,50)} = 0.2353$, p=0.6297) **B.** GlcCer C20:0 (*Interaction*: $F_{(3,48)}= 0.6806$, p=0.5682, *Age*: $F_{(3,48)} = 0.7990$, p=0.5005, and *Genotype*: $F_{(1,48)} =1.619$, p=0.2094) **C**. GlcCer C22:0 (*Interaction*: $F_{(3,50)}= 0.3007$, p=0.8247, *Age*: $F_{(3,50)} = 1.027$, p=0.3886, and *Genotype*: $F_{(1,50)} = 2.898$, p=0.0949) and **D.** GlcCer C24:1 (*Interaction*: $F_{(3,50)}= 0.7606$, p=0.5215, *Age*: $F_{(3,50)} = 1.821$, p=0.1554, *Genotype*: $F_{(1,50)} = 9.377$, ***p=0.0035) were examined in the striatum of the same mice groups and time points. Error bars represent SEM. For all lipids, 3mo $GBA1^{+/+}$ N= 8, 3mo $GBA1^{+/L444P}$ N=7 except C18:0 N=5 and C20:0 N=6, 6mo $GBA1^{+/+}$ N= 8 except C18=10 and C20 =7, 6mo $GBA1^{+/L444P}$ N=8, 9mo $GBA1^{+/+}$ N= 6, 9mo $GBA1^{+/L444P}$ N=7, 12mo $GBA1^{+/+}$ N= 7, and 12mo $GBA1^{+/L444P}$ N=7.*p<0.05, **p<0.01, ***p<0.001, and ****<0.0001.

### Extended Figure 5-1

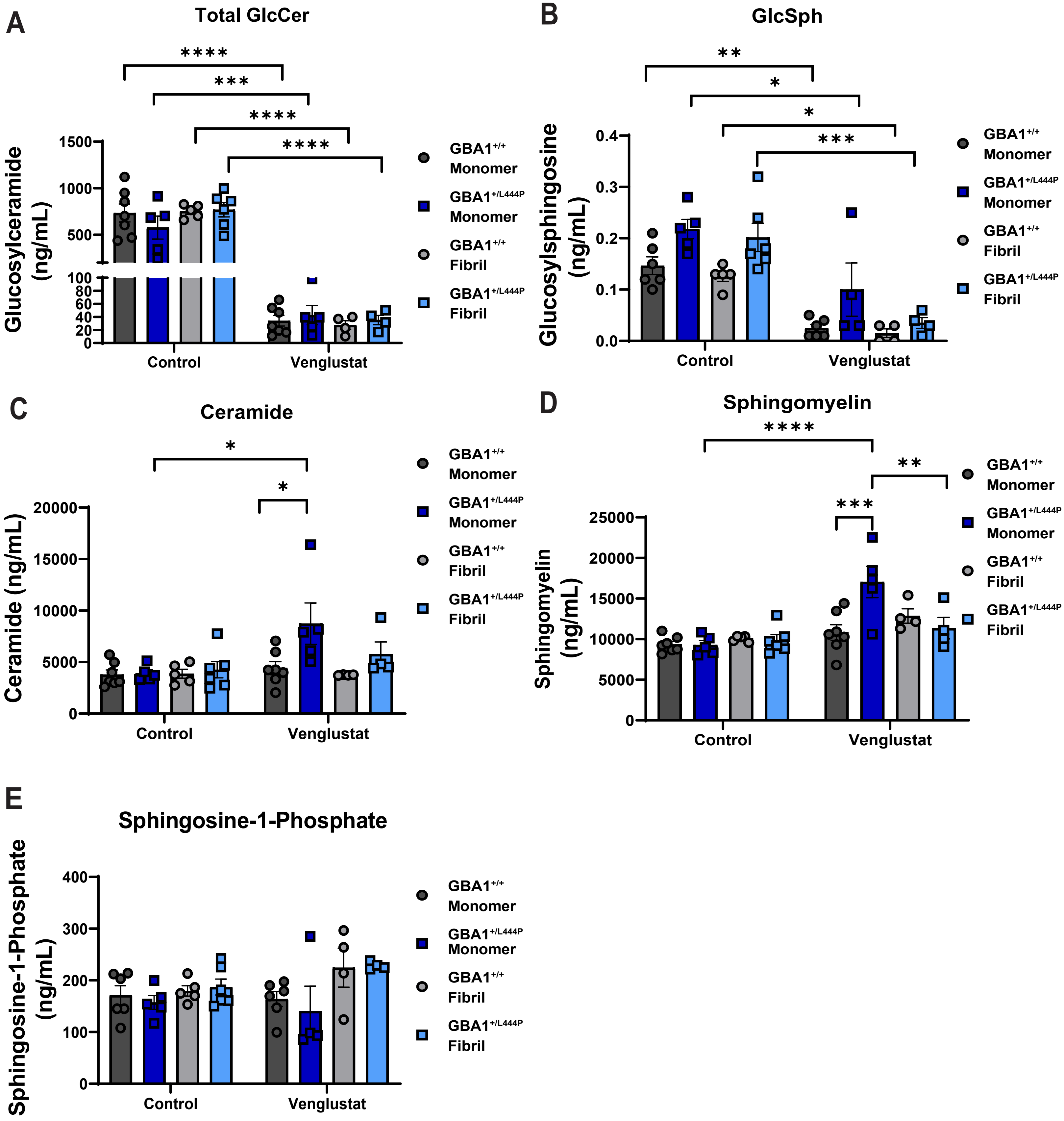

**Extended Data 5-1: Evaluation of lipid plasma concentrations.** Plasma was collected from $GBA1^{+/+}$ and $GBA1^{+/L444P}$ at 10-months post-monomer or fibril injection. Mass Spectrometry was used to quantify **A**. Total GlcCer levels (Three-way ANOVA; *Interaction*: $F_{(1,35)}$= 0.7606, p=0.3891, *Drug Treatment x α-syn Treatment:* $F_{(1,35)}$= 1.268, p=0.2677, *Drug Treatment x Genotype*: $F_{(1,35)}$= 0.6130, p=0.4389, *α-syn Treatment x Genotype:* $F_{(1,35)}$= 0.7431, p=0.3945, *Drug Treatment*: $F_{(1,35)}$= 181.7, ****P<0.0001, *α-syn Treatment*: $F_{(1,35)}$= 0.9673, p=0.3321, *Genotype*: $F_{(1,35)}$= 0.3836, p=5394), **B**. GlcSph (*Interaction*: $F_{(1,35)}$= 0.8940, p=0.3515, *Drug Treatment x α-syn Treatment:* $F_{(1,35)}$= 0.3667, p=0.5491, *Drug Treatment x Genotype*: $F_{(1,35)}$= 0.6867, p=0.4134, *α-syn Treatment x Genotype:* $F_{(1,35)}$= 0.6519, p=0.4254, *Drug Treatment*: $F_{(1,35)}$= 67.97, ****p<0.0001, *α-syn Treatment*: $F_{(1,35)}$= 3.186, p=0.0838, *Genotype*: $F_{(1,35)}$= 14.87, ***p=0.0005). **C.** Ceramide (*Interaction*: $F_{(1,35)}$= 0.9955, p=0.3253, *Drug Treatment x α-syn Treatment:* $F_{(1,35)}$= 2.536, p=0.1203, *Drug Treatment x Genotype*: $F_{(1,35)}$= 5.255, *p=0.0280, *α-syn Treatment x Genotype:* $F_{(1,35)}$= 0.6352, p=0.4308, *Drug Treatment*: $F_{(1,35)}$= 7.185, *p=0.0111, *α-syn Treatment*: $F_{(1,35)}$= 1.582, p=0.2168, *Genotype*: $F_{(1,35)}$= 7.241, *p=0.0109) **D.** Sphingomyelin (*Interaction*: $F_{(1,35)}$= 7.734, **p=0.0087, *Drug Treatment x α-syn Treatment:* $F_{(1,35)}$= 3.327, p=0.0767, *Drug Treatment x Genotype*: $F_{(1,35)}$= 3.289, p=0.0783, *α-syn Treatment x Genotype:* $F_{(1,35)}$= 8.184, **p=0.0071, *Drug Treatment*: $F_{(1,35)}$= 24.29, ****p<0.0001, *α-syn Treatment*: $F_{(1,35)}$= 0.7393, p=0.3957, *Genotype*: $F_{(1,35)}$= 2.626, p=1141), and **E**. Sphingosine 1-Phosphate (*Interaction*: $F_{(1,32)}$= 0.008874, p=0.9255, *Drug Treatment x α-syn Treatment:* $F_{(1,32)}$= 3.188, p=0.0836, *Drug Treatment x Genotype*: $F_{(1,32)}$= 0.04317, p=0.8367, *α-syn Treatment x Genotype:* $F_{(1,32)}$= 0.6178, p=0.4376, *Drug Treatment*: $F_{(1,32)}$= 1.016, p=0.3209, *α-syn Treatment*: $F_{(1,32)}$= 9.109, **p=0.0050, *Genotype*: $F_{(1,32)}$= 0.1771, p=0.6767). GlcCer, ceramide, and sphingomyelin had $GBA1^{+/+}$ N=7, $GBA1^{+/L444P}$ N=5, fibril treated $GBA1^{+/+}$ N=5, and fibril treated $GBA1^{+/L444P}$ N=6 for control treated groups and $GBA1^{+/+}$ N=7, $GBA1^{+/L444P}$ N=5, fibril treated $GBA1^{+/+}$ and $GBA1^{+/L444P}$ N=4 for venglustat treated groups. GlcSph and sphingosine 1-Phosphate had $GBA1^{+/+}$ N=6, $GBA1^{+/L444P}$ N=4, fibril treated $GBA1^{+/+}$ and $GBA1^{+/L444P}$ N=4 venglustat treated mice. For GlcSph and sphingosine 1-phosphate, $GBA1^{+/+}$ N=6, $GBA1^{+/L444P}$ N=5, and fibril treated $GBA1^{+/+}$ N=5. GlcSph had fibril treated $GBA1^{+/L444P}$ mice N=5, whereas sphingosine 1-phosphate had N=6 for the same group. *p<0.05, **p<0.01, ***p<0.001, and ****<0.0001.

Extended Figure 5-2

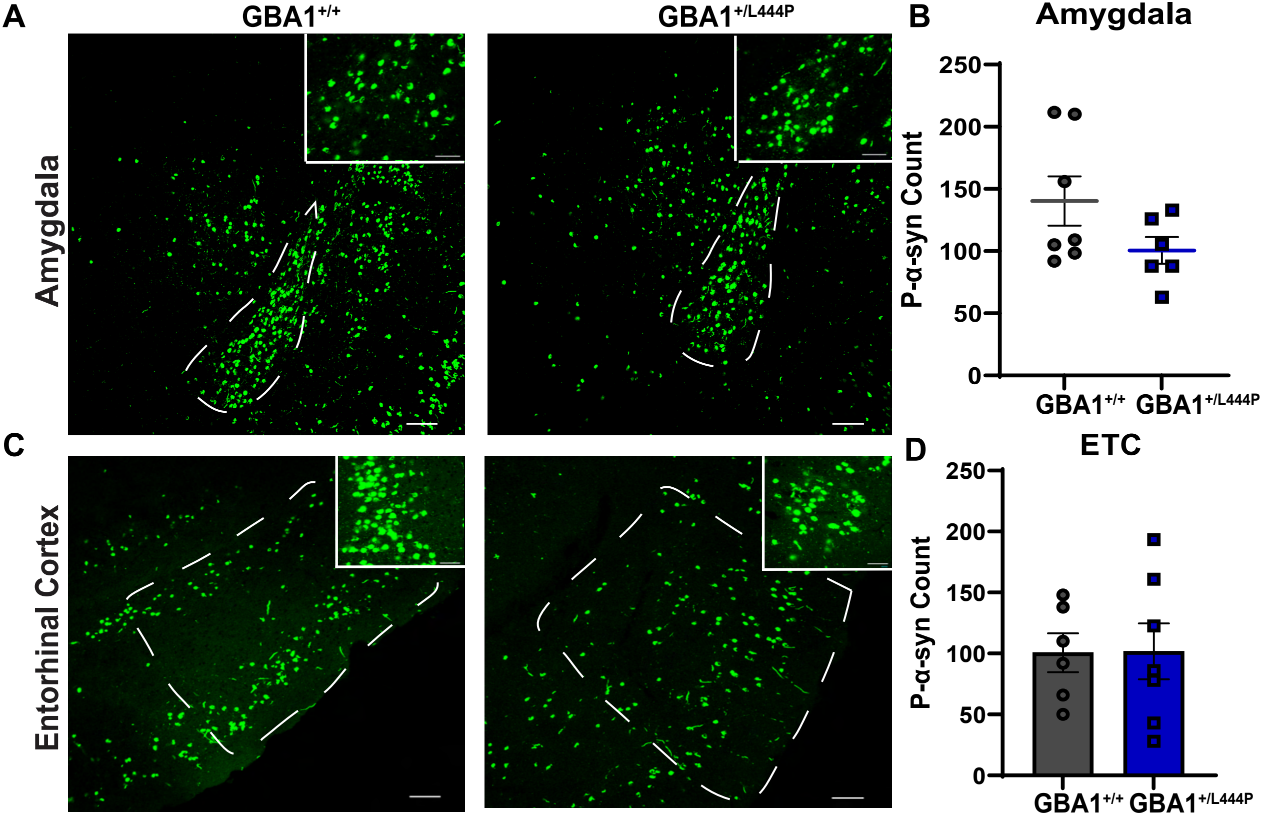

**Extended Figure 5-2: Aggregate burden in the entorhinal cortex and basolateral amygdala. A.** Representative images of $GBA1^{+/+}$ and $GBA1^{+/L444P}$ fibril-injected mice of the amygdala were captured using confocal microscopy. Scale bar =100um, zoomed photos scale bar = 50um) **B**. Quantification of p-α-syn in the amygdala (N=7 for $GBA1^{+/+}$, N=6 for $GBA1^{+/L444P}$, Mann-Whitney test: W= 11, p=0.1690). The amygdala aggregate data failed the test for normality and graphed as a scatter plot without a bar using a Mann-Whitney test. Error bars represent SEM. **C**. Representative images of $GBA1^{+/+}$ and $GBA1^{+/L444P}$ fibril-injected mice of the entorhinal cortex were captured using confocal microscopy. Scale bar =100um, zoomed photos scale bar = 50um. **D.** Quantification of p-α-syn in the entorhinal cortex (N=6 for $GBA1^{+/+}$, and N=7 $GBA1^{+/L444P}$: Independent t-test: $t_{(11)} = 0.03628$; p=0.9717). Error bars represent SEM.

**Extended Figure 5-3**

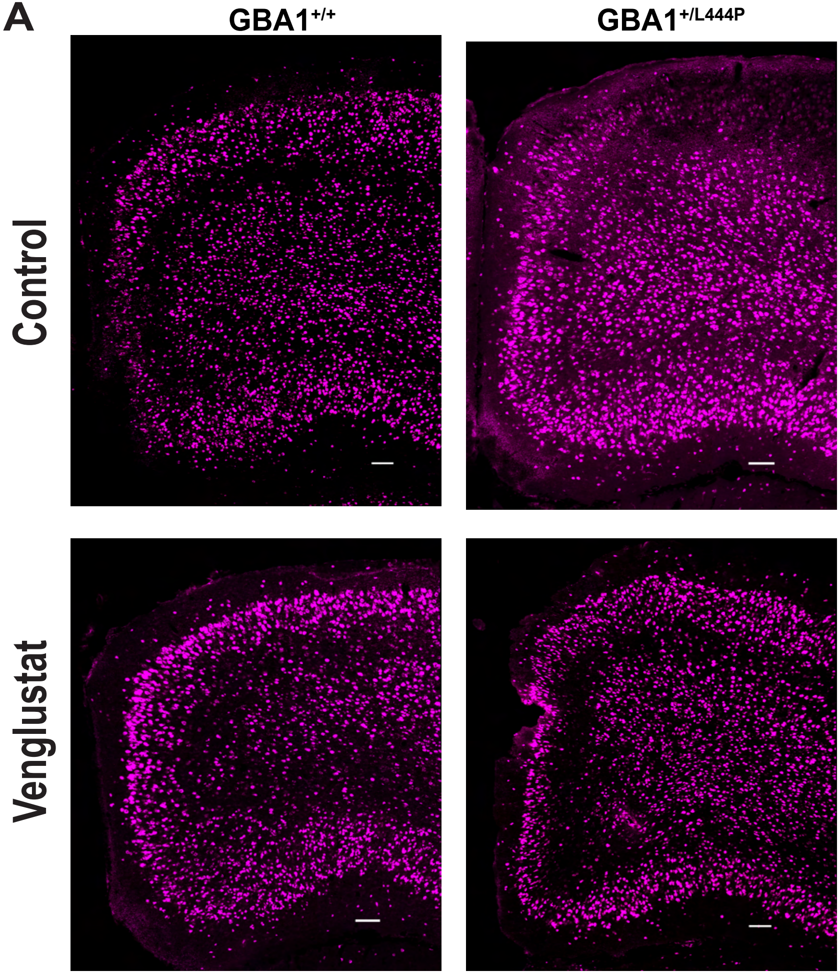


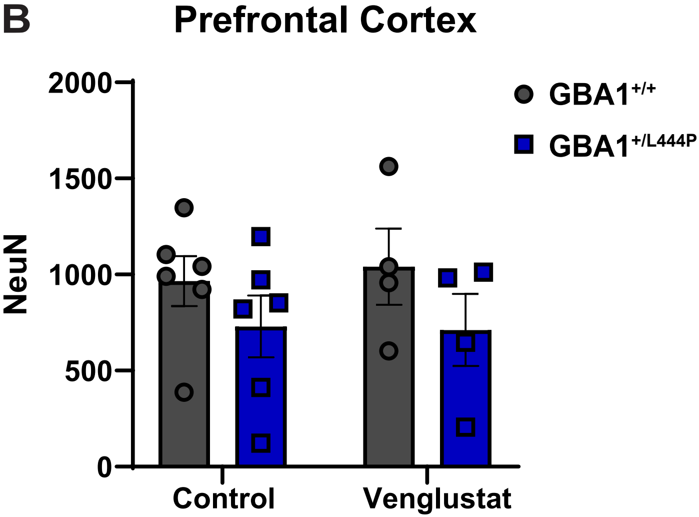

**Extended figure 5-3**: **NeuN analysis of the Prefrontal Cortex.** **A**. 10-months after bilateral striatal injection, immunofluorescence for NeuN was performed. Representative images of $GBA1^{+/+}$ and $GBA1^{+/L444P}$ fibril-injected mice fed control or venglustat chow of the dmPFC were captured using confocal microscopy. Scale bar =100um. **B**. Quantification of NeuN in the dmPFC (N=6 for both control chow groups, N=4 for venglustat chow groups, Two-way ANOVA: *Interaction*: $F_{(1,16)}$= 0.07678, p=0.7853, *Drug Treatment*: $F_{(1,16)}$ = 0.02827, p=0.8686, *Genotype*: $F_{(1,16)}$= 2.821, p=0.1124). Error bars represent SEM.

**Extended Figure 5-4**

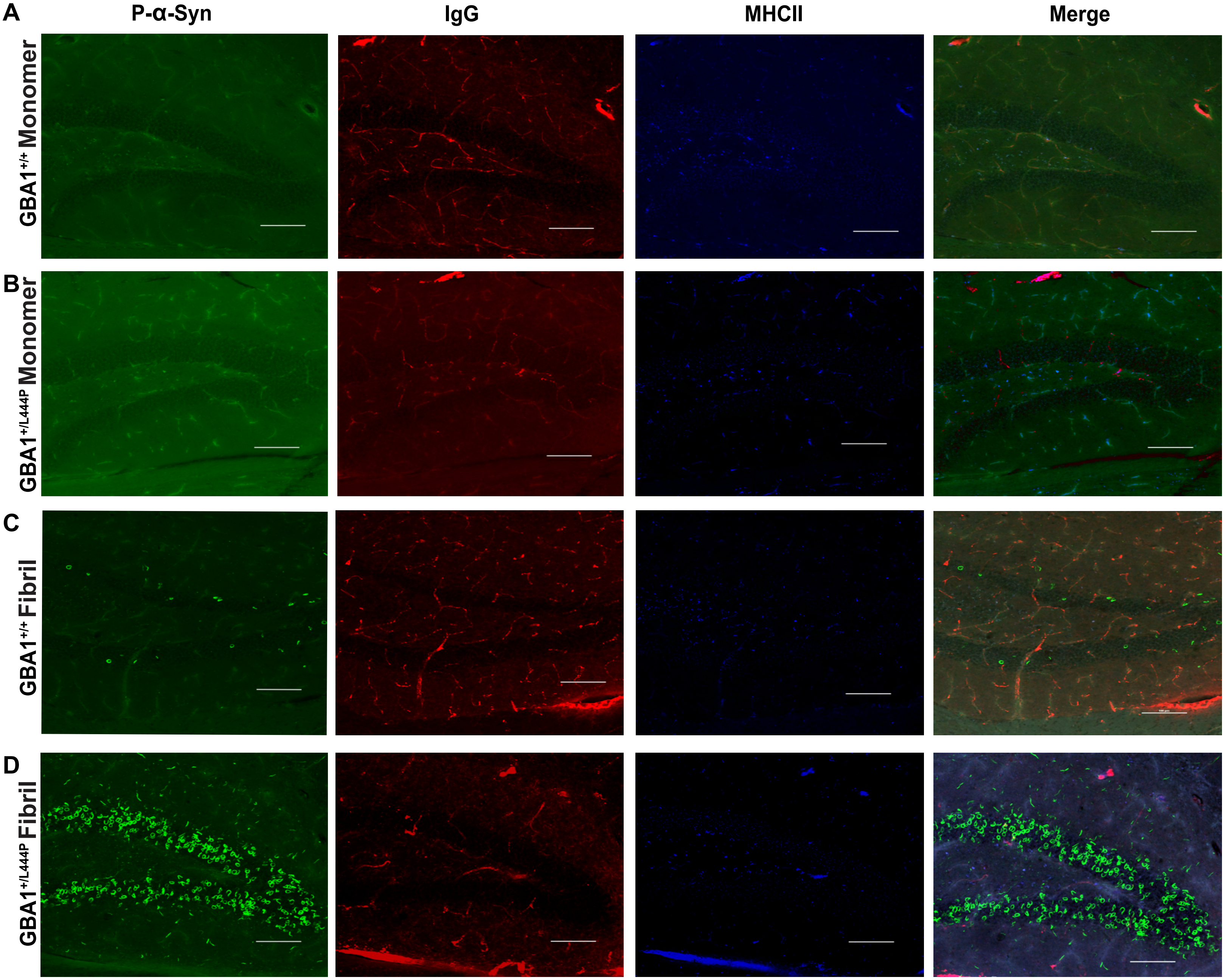

**Extended Figure 5-4: Neuroinflammation of the Granule Cell layer of the dentate gyrus of the hippocampus.** 10-months after bilateral striatal injection, immunofluorescence for p-α-syn, IgG, and MHCII was performed. Representative images of the granule cell layer of the dentate gyrus of the hippocampus of **A**. monomer-injected GBA1$^{+/+}$, **B**. monomer-injected GBA1$^{+/L444P}$, **C**. fibril-injected GBA1$^{+/+}$, and **D**. fibril-injected GBA1$^{+/L444P}$ mice were captured using confocal microscopy. Scale bar =100um. N=3 for all groups.
